# Tumor-educated monocyte-dendritic progenitors promote a metastatic switch

**DOI:** 10.1101/2020.08.25.266189

**Authors:** Ksenia Magidey-Klein, Ksenya Kveler, Tim J. Cooper, Rachelly Normand, Tongwu Zhang, Michael Timaner, Ziv Raviv, Brian James, Roi Gazit, Ze’ev A. Ronai, Shai S. Shen-Orr, Yuval Shaked

**Affiliations:** Department of Cell Biology and Cancer Science, Rappaport Faculty of Medicine, Technion Integrated Cancer Center, Technion, Haifa, Israel; Faculty of Medicine, Technion-Israel Institute of Technology, Haifa, Israel; Division of Cancer Epidemiology and Genetics, Laboratory of Translational Genomics, National Cancer Institute, Bethesda, MD; Sanford Burnham Prebys Medical Discovery Institute, San Diego, CA; National Institute for Biotechnology in the Negev, Ben-Gurion University of the Negev, Beer-Sheva, Israel; Faculty of Biology, Technion-lsrael Institute of Technology, Haifa, Israel

## Abstract

Myeloid skewing of hematopoietic cells is a prominent promoter of metastasis. However, little is known about their education and differentiation pattern from hematopoietic stem and progenitor cells (HSPCs) during tumor progression and metastasis. Here we show that metastatic tumors dictate a unique differentiation pattern of HSPCs towards a specific myeloid progeny. Using single cell RNA-sequencing analysis integrated with proteomic screen of tumor secretome, we demonstrate that highly metastatic tumors dictate a long-lived differentiation of HSPCs towards monocyte-dendritic progenitors (MDPs) while low-metastatic tumors promote their differentiation into granulocyte-monocyte progenitors (GMPs). This effect is driven by IL-6 axis that is highly active in metastatic tumors. Consequently, loss and gain of function of IL-6 in tumor cells resulted in decreased and increased metastasis and corresponding MDP levels, respectively. Consistently, IL-6-educated MDPs but not GMPs obtained from highly metastatic tumors, adoptively transferred into mice bearing low metastatic tumors resulted in increased metastasis due to their further differentiation into immunosuppressive (M2) macrophages. Overall, our study reveals a new role for tumor-derived IL-6 that hijacks HSPC differentiation program towards myeloid cells that contribute to metastasis.

## Introduction

Despite recent advances in cancer therapies, metastasis remain mostly incurable, accounting for the majority of cancer-related mortalities. Metastasis is a multistep process that involves dissemination of cancer cells from the primary tumor mass, intravasation into blood circulation, seeding and subsequent growth at distant sites (Mehlen and Puisieux, 2006). Successful colonization of tumor cells at distant sites relies, in part, on bone marrow derived cells (BMDCs) that are recruited earlier than tumor cells to the distant sites where they contribute to the formation of a pre-metastatic niche (Kaplan et al., 2005; Wang et al., 2019). Among these cells are immunosuppressive myeloid cells such as M2 macrophages and myeloid derived suppressor cells (MDSCs) (Doak et al., 2018; Gabrilovich, 2017).

The immune cells within tumors all originate from HSPCs. HSPCs are sensitive to external stimuli, such as infection or injury. In response, they enter an extensive proliferative phase, in order to increase the reservoir of required immune cells, shifting the balance between various populations of leukocytes. Once the stimulus ceases, HSPCs return to a quiescent stage, maintaining a normal homeostasis (Hoggatt et al., 2016). Tumors also secrete external stimulus which contribute to HSPC differentiation. It is well established that tumors induce myelopoiesis as a means of supplying immature progenitors that act as immunosuppressive cells. A recent study demonstrated that the composition of circulating HSPCs is significantly altered in patients with solid tumors, with increased levels of granulocyte-monocyte progenitors (GMPs) and a general bias towards granulocyte formation (Wu et al., 2014). In addition, elevated numbers of circulating HSPCs correlate with tumor progression and decreased overall survival (Wildes et al., 2019; Wu et al., 2014). Moreover, it has been suggested that tumor-mediated signals directly affect the fate of HSPCs, leading to their differentiation into MDSCs that help forming a premetastatic niche (Gabrilovich et al., 2012; Wang et al., 2019). However, the specific interactions between highly metastatic tumor cells and HSPC differentiation to promote metastasis, have not been elucidated.

Here we show that highly metastatic tumors regulate HSPC fate, shifting their differentiation pattern towards tumor and metastasis supporting cells. HSPC differentiation induced by highly metastatic tumors, is dictated by IL-6 signaling. This crosstalk leads to increased levels of monocyte-dendritic progenitors (MDPs) that further give rise to a immunosuppressive macrophage phenotype, ultimately promoting metastasis.

## Methods

### Tumor cell cultures

The 4T1 murine breast carcinoma cell line and B16-F1 and B16-F10 murine melanoma cell lines were purchased from the American Type Culture Collection (ATCC, Manassas, VA, USA). Cells were used within 6 months of resuscitation. 67NR murine breast carcinoma cells were kindly provided by Prof. Sleeman (Menheim, Germany). 4T1 and 67NR cell lines have the same genetic background, and represent cells with high and low metastatic potential, respectively, as previously described (Schmaus and Sleeman, 2015). The cells were grown in Dulbecco’s modified Eagle’s medium (DMEM, SigmaAldrich, Rehovot, Israel). B16-F10 and B16-F1 cells represent cells with high and low metastatic potential, respectively, as previously described (Ishiguro et al., 1996). For overexpression of IL-6, B16-F1 cells were transfected with 1 μg DNA of pCMV3 vector encoding IL-6 or the empty control pCMV3 vector (EV) using PolyJet (SignaGen Laboratories, MD, USA) following the manufacturer’s instructions. For generating stable clones, transfected cells were grown for two weeks with hygromycin (250μg/ml). Conditioned medium obtained after 24 hours was analyzed for IL-6 levels using ELISA. Cell proliferation of IL-6 overexpressing cells was carried out by XTT, as previously described (Chang et al., 2020). Cells were grown in Roswell Park Memorial Institute medium (RPMI, SigmaAldrich) and supplemented with 1.5% non-essential amino acids, 0.75 % MEM vitamins and 1.5 % sodium bicarbonate. All cell media were supplemented with 5% fetal calf serum (FCS), 1% L-glutamine, 1% sodium pyruvate, and 1% Pen-Strep-Neomycin in solution (purchased from Biological Industries, Israel). The cells were cultured in a humidified chamber in 5% CO_2_ at 37°C. Cells were routinely tested for mycoplasma and found to be mycoplasma-free.

### Animals

Female BALB/c and C57BL/6 mice (8-10 weeks of age) were purchased from Envigo (Israel) or Jackson laboratories (Bar Harbor ME, USA). For bone marrow transplantation, co-inoculation and adoptive transfer experiments, transgenic C57BL/6 mice expressing enhanced-GFP under the regulation of the ubiquitin promoter were used (B6-EGFP, The Jackson Laboratory, Bar Harbor, ME). The Animal Care and Use Committees of the Technion (Haifa, Israel) and Sanford Burnham Prebys (San Diego, CA) approved all animal studies and experimental protocols.

### Murine tumor models

4T1 and 67NR cells (5×10^5^ cells in 50 μl serum free medium) were orthotopically implanted in the mammary fat pad of BALB/c female mice. B16-F1 and B16-F10 cells (5×10^5^ cells in 200 μl serum free medium) were subdermally implanted into the flanks of C57BL/6 mice. Tumor volume was measured twice a week with Vernier calipers and calculated according to the formula width^2^xheightx0.5. When tumor size reached approximately 1500 mm^3^ (endpoint), mice were sacrificed and tumor, lungs and femurs were removed. Blood was drawn by cardiac puncture prior to mice euthanasia.

In some experiments, mice were treated with anti-IL-6 antibodies (20mg/kg, MP5-20F3 clone, BioXCell, NH, USA) or IgG control antibodies (20mg/kg, Clone MP5-20F3), twice weekly, as previously described ^12^. Twenty-four hours after the last injection, the mice were sacrificed for further pathological and cellular analysis.

In co-inoculation experiments, Lineage negative (Lin-) cells (100,000 cells/mouse) or Lin-Sca1+Kit+ (LSK) cells (25,000 cells/mouse) obtained from bone marrow of tumor-free GFP-expressing mice were mixed with B16-F1 or B16-F10 cells in Matrigel (BD Biosciences, CA, USA) and subdermally implanted into the flanks of C57BL/6 mice. Lin- and LSK cells mixed with Matrigel alone were used as control groups in order to evaluate spontaneous differentiation in Matrigel plugs. Lin-cells were purified using the MagCellect Mouse Hematopoietic Cell Lineage Depletion Kit (R&D, MN, USA). LSK cells were sorted using FACSAria™ IIIu (BD Biosciences, CA, USA). Mice were sacrificed 14 days (for Lin-) and 10 days (for LSK) after implantation, and tumors were removed.

### Flow cytometry acquisition and analysis

Lungs and tumors were prepared as single cell suspensions as previously described ^13^. Bone marrow cells were flushed from the bone marrow and peripheral blood was collected either by retro-orbital bleed or cardiac puncture. Cells were immunostained with specific antibodies against surface markers to define various cell types listed in Table S1. Conjugated monoclonal antibodies were purchased from BD Biosciences (San Jose, CA) or BioLegend (San Diego, CA). Sca1(D7)-PE/BV786, CD117(2B8)-APC, CD34(HM34)-PE, IL-7R(A7R34)-PE-Cy7, Flt3(A2F10)-PE-Cy5, F4/80(BM8)-PE, CD11b(M1/70)-PerCP, Gr-1(RB6 8C5)-BV510, FCγR(93)-BV510, CD115(AFS98)-PeCy7, CD206(C068C2)-BV421, CD11c(N418)-APC-Cy7, Ly6C(1A8)-BV605, Ly6G(HK1.4)-BV510, CD3ε(30-F11)-Alexa Fluor 700, B220(RA3-6B2)-BV605 and lineage cocktail (17A2/RB6-8C5/RA3-6B2/Ter 119/M1/70)-Alexa Fluor 700 or BV421. For phospho-STAT3 (p-STAT3) analysis, naïve bone marrow cells were stimulated first with escalating doses of IL-6 for 30 minutes at 37°C, and subsequently were immunostained for MDPs. Right after, the cells were fixed in 1.6% paraformaldehyde and permeabilized in ice-cold 90% ethanol. Cells were immunostained with p-STAT3 (Tyr705)-AF488 at 4 °C for 40 minutes. At least 500,000 events were acquired for each sample using BD LSRFortessa cytometer and analyzed with FlowJo 10.2 software.

### Bone marrow transplantation

Donor GFP-expressing mice were inoculated with either B16-F1 or B16-F10 tumor cells. At endpoint, the mice were sacrificed and bone marrow cells were obtained by flushing the femur and tibia with PBS. Lineage depletion kit (R&D, MN) was used to purify immature cells. Enriched cells were immunostained and sorted for Lin-or LSK (1×10^5^ and 5×10^3^ cells, respectively), and subsequently intravenously transplanted into recipient lethally irradiated mice (10 Gy in 8 min, single dose) together with supportive total bone marrow cells (1×10^6^ cells/mouse), obtained from naïve (non-GFP) C57BL/6 mice. Recipient mice exhibiting less than 1% total chimerism were considered as failed transplantations and excluded from the analysis. Reconstitution recovery of GFP+ donor cells was tested following 16 weeks, by collecting the blood from the retro-orbital venous plexus.

### MDP Adoptive transfer

GFP-expressing mice were implanted with B16-F10 tumor cells, and then treated with anti-IL-6 or IgG control antibodies, as described above. Lin-cells were purified from the bone marrow using a lineage depletion kit (R&D, MN) and immunostained for MDP sorting. MDPs (1×10^3^ cells) were intravenously injected to naive C57BL/6 mice, which were subsequently implanted with B16-F1 tumor cells. When the tumors reached endpoint, the mice were sacrificed and MDP differentiation was evaluated by analyzing the GFP+ cells using flow cytometry. Lungs were harvested and subjected to histopathological staining for metastasis scoring.

### Mass cytometry acquisition and analysis

High throughput mass-cytometry (CyTOF) analysis was performed as previously described (Shaked et al., 2016). Briefly, tumors were prepared as single cell suspensions. Cells were pooled (3×10^6^) and immunostained with a mixture of metal-tagged antibodies using the different surface markers as indicated in Table S2. All antibodies were conjugated using the MAXPAR reagent (Fluidigm, CA, USA) and tittered prior to staining. Rhodium and iridium intercalators were used to identify live/dead cells. Cells were washed twice with PBS, fixed in 1.6% formaldehyde (SigmaAldrich, MO, USA), washed again in ultrapure H_2_O, and acquired by CyTOF mass cytometry system (Fluidigm, CA, USA). Acquired data was uploaded to Cytobank web server (Cytobank Inc.). CD11b+ myeloid live cells were used for the analysis, and the gated cells were segregated into sub-population clusters by expression markers. Data analysis was performed by viSNE algorithm (Amir el et al., 2013), via the Cytobank server. Changes in specific populations were validated by flow cytometry.

### Colony forming assay

For mouse colony forming units (CFU), red blood cells were lysed from peripheral blood of tumor-free or tumor-bearing mice. The cells were then seeded in triplicates at a concentration of 100,000 cells/well into 6-well culture plates with M3434 methylcellulose (Methocult, Stem Cell Technologies, Vancouver, Canada). For assessing the differentiation pattern of naïve bone marrow cells and MDPs in the presence of tumor-derived conditioned medium (TCM), incomplete methylcellulose medium M3431 (Stem Cell Technologies, Vancouver, Canada) was supplemented with 30% TCM obtained from 5×10^6^ total tumor cells after they were adjusted to culture for 24 h in serum free medium. BM cells (10,000 cells/well) and sorted MDP cells (50 cells/well) were plated for 12 days. Plates were imaged with a Zeiss microscope and colonies were scored.

### Single cell RNA sequencing and data analysis

LSK cells were sorted from bone marrow of met-low and met-high melanoma tumor bearing mice based on expression markers defined in Table S1. Briefly, LSK cells were sorted into two 384-well plates (Biorad Laboratories), one for each group, containing RT mix, and barcoded 3’ RT primer, in nuclease-free water. The plates were kept at −80°C until analyzed. Single cell RNA sequencing (scRNA-seq) was performed at the New York University Genome Center. On the day of library preparation, plates were removed from −80°C and placed in the thermal cycler for thaw, lyse and annealing purposes (Hold for 22°C, 22C for 2 min, 72°C for 3 min, Hold at 4°C). After the program ended, 2 μl of RT Mix 2 [each reaction consisting of 0.5 μl 5’ Custom TSO Primer 10 μM (IDT Technologies), 0.925 μl 5M Betaine (ThermoFisher Scientific), 0.4 μl 100 mM MgCl_2_ (Sigma Aldrich,CA,USA), 0.125 μl Superase In RNase Inhibitor 20 U/ μl (Invitrogen), and 0.05 μl Maxima H Minus RT enzyme (200 U/ μl)] was added to each well and samples were sealed and then mixed using the Eppendorf thermomixer at 2000 RPM for 30 seconds at room temperature. The plates were briefly centrifuged at 2000 RPM for 30 seconds at 4°C. Plates were then placed in the thermal cycler for the Maxima_RT program (Hold at 50°C, 50°C for 94 minutes, 85°C for 5 minutes, Hold at 4°C). After the RT program, each sample was treated with 7 μl cDNA PCR Master mix [each reaction consisting of 0.25 μl 10 μM IS-PCR Primer Mix (IDT Technologies), 0.5 μl molecular grade water, and 6.25 μl 2X KAPA Hifi ReadyMix (Roche)]. Samples were mixed on the thermomixer at 2000 RPM for 30 seconds at RT, and centrifuged briefly at 2000 RPM for 30 seconds at 4°C. Plates were placed in the thermal cycler for 3-Step cDNA amplification (Hold at 98°C, 13 cycles of [98° C for 15 sec; 75°C for 20 sec; 72°C for 6 min], 5 cycles of [98°C for 15 sec; 72°C for 20 sec; 72°C for 6 min], 13 cycles of [98°C for 15 sec; 67°C for 20 sec; 72°C for 6 min], 72°C for 5 minutes, Hold at 4°C). After cDNA amplification, samples were stored at 4°C until ready for cDNA pooling and cleanup. The pools were first cleaned with a 0.7X SPRISelect bead cleanup followed by a 0.6X SPRISelect bead cleanup and eluted in 25 μl elution buffer. Pooled cDNA was quantified with Qubit HS DNA (Invitrogen) and Fragment Analyzer HS-NGS (Agilent). Pooled and purified cDNA (600 pg) was taken into Nextera XT (Illumina) library prep and tagmented according to the manufacturer’s instructions. The samples were given custom P7 and P5 adapters (10 uM, 1 μl for each adapter) and amplified using a modified NXT scRNA PCR program (Hold at 95°C, 95°C for 30 seconds, 13 cycles of [95°C for 10 seconds, 55°C for 30 seconds, 72°C for 30 seconds], 72°C for 5 minutes, Hold at 4°C). Each final library was quantified with Qubit HS DNA (Invitrogen) and Fragment Analyzer HS-NGS (Agilent), and then diluted to 2 nM. The libraries were loaded at a concentration of 8.5 pM on a HiSeq Rapid Run with run cycles of 26×9×9×76 and were compiled into a digital gene expression matrix used for downstream analyses.

For data filtering and clustering, Seurat R package v3.00 (Butler et al., 2018) was used. Cells with less than 1000 non-zero genes were filtered out; genes expressed in less than 3 cells and non-coding genes were omitted. Outlier cells with more than 5% of expressed genes coming from mitochondria were removed from the analysis, as high mitochondrial expression likely indicates cells undergoing apoptosis. Altogether, the filtered data contained 758 cells and 13,997 genes. Following log-normalization, the top 2000 variable genes were identified and used to find Louvain clusters using the default resolution of 0.8, as well as resolution set to 1 to obtain a larger number of communities. Top 75 significant principal components as selected with the JackStraw method, were and used to compute uniform manifold approximation and projection (UMAP). To annotate individual cells with main cell types, we applied the SingleR package v0.22 (Aran et al., 2019), using ImmGen gene expression compendium (Heng et al., 2008) as the reference dataset. This resulted in the most likely cell type being inferred for each single cell independently, in an unsupervised manner, based on the expression profile of the cell. Annotation significance p-values were computed using SingleR built-in chi-squared outlier test for the top-scored cell type match. The resulting cellular origins were used to calculate cell type proportions in bone marrow and compare them between met-high and met-low mice. For determining proportion difference significance, the bias-corrected and accelerated (BCa) bootstrap method for independent two-samples was applied, together with one-sided hypothesis testing, using wBoot R Package v1.0.3. Differential gene expression analysis and gene set enrichment analysis (GSEA) was performed using MAST (Finak et al., 2015), a package suited to analysis of sparse scRNA-seq, and clusterProfiler (Yu et al., 2012), respectively. GSEA reference signatures were obtained from MSIGDB (Liberzon et al., 2015) (v7.1 - Hallmark (H), Regulatory Target Gene Sets (C3)).

### Cytokine array and ELISA

Tumor derived conditioned medium (TCM) was obtained from 5×10^6^ total tumor cells that underwent single cell suspension and subsequently cultured for 24 h in serum-free medium. TCM from met-low or met-high B16 melanoma tumors were applied to a proteome profiler mouse XL cytokine array (ARY028, R&D, MN) in accordance with the manufacturer’s instruction. The signals corresponding to each factor in the array were quantified by densitometry analysis. The ratio between the levels of the various factors in met-high and met-low TCM was calculated. IL-6 levels in TCM of met-low and met-high melanoma and breast carcinoma tumors were quantified using a specific enzyme-linked immunosorbent assay (ELISA) kit (ab46100, Abcam, US). The ELISA experiments were performed with 5-8 mice per group, and analyzed as mean ± SD.

### NanoString platform

For NanoString gene expression analysis, the gene panel nCounter PanCancer Progression was used (catalog #XT-CSO-PROG1–12), which contains probes for 740 test genes and 30 housekeeping genes. RNA was purified with Single Cell RNA Purification kit (Norgen, Canada) from FACS sorted CD45+ live cells and analyzed with the nCounter platform. Data was filtered and normalized using a generalized linear model (GLM) (R package, NanoStringDiff (v. 1.16.0))(Wang et al., 2016) - based on positive controls, housekeeping genes and estimated background. Differential gene expression analysis was performed (met-high vs. met-low) using either (i) a negative binomial model, akin to DESeq2 (Love et al., 2014) and (ii) AUC-ROC curves (R package, Seurat (v. 3.1.4))(Butler et al., 2018)- collectively revealing 7 significant DEGs (FDR p-value < 0.05, fold-change > 1.25, AUC = 1). An AUC value of 1 denotes the ability of a given gene to perfectly distinguish met-high from met-low.

### Multi-omics data analysis for characterization of the crosstalk between tumor cells and BM

To identify the axis responsible for the crosstalk between tumor cells and the BM compartment, differential expression of cytokine receptor genes in the scRNA-seq BM data was analyzed and integrated with the tumor cytokine array findings. First, following SingleR cell type annotation of individual cells, the Wilcoxon rank-sum test method using the Seurat R package was applied, to identify, per cell type, the genes differentially expressed in met-high versus met-low groups. The testing was limited to genes showing, on average, at least 1.2-fold difference and those detected in at least 10% of cells in either of the groups. Multiple comparison adjustment was carried out using Benjamini–Hochberg correction. Next, cytokines in the array were paired to their respective receptors within the differentially expressed genes, based on the KEGG cytokine–cytokine receptor interaction pathway entry (Kanehisa et al., 2016). This resulted in the list of cytokine-receptor pairs, per cell type, ranked by their cytokine expression ratio between the groups, receptor fold-change and its adjusted p-value of differential expression.

### Statistical analysis for in vitro and in vivo studies

Data are expressed as mean ± standard deviation (SD). For in vitro studies, to ensure adequate statistical power, all experiments were performed with two technical repeats and at least three biological repeats. In the in vivo studies, mice that demonstrated pathological symptoms not related to their condition or disease were excluded from the study and the analysis. Mice were randomized before tumor implantation. The analysis of the results was performed blindly. At least 5 mice per group were used in order to reach statistical power considering a Gaussian distribution. The statistical significance of differences was assessed by one-way ANOVA, followed by Turkey post hoc statistical test when comparing between more than two sets of data or by two-tailed student’s t-test when comparing between two sets of data, using GraphPad Prism 5 software (La Jolla, CA). Significance was set at values of p⍰<⍰0.05.

## Results

### Metastatic tumors promote hematopoiesis and myeloid skewing

To study the communication occurring between BM niche and metastatic tumor cells, we used tumor cell line pairs comprising one line that rarely metastasizes (met-low) and another line that metastasizes with high frequency (met-high). Cell lines within each pair originated from the same parental cell line, allowing a biologically relevant comparison. Mice were either sub-dermally implanted in the flank with met-low B16-F1 or met-high B16-F10 melanoma cells, or implanted in the mammary fat pad with met-low 67NR or met-high 4T1 breast carcinoma cells. In both tumor models, met-low and met-high primary tumors grew at similar rates, whereas lung metastasis was significantly increased in the met-high groups (Fig. S1A-C), in agreement with previous studies (Ishiguro et al., 1996; Schmaus and Sleeman, 2015). At endpoint, HSPCs in the BM compartment of naïve and tumor-bearing mice were analyzed by flow cytometry. Early HSPCs were defined as Lin-, Sca1+ and CD117+, representing LSK cells, as previously described (van de Rijn et al., 1989). The percentage of LSK in the BM of mice implanted with met-high tumors was increased compared to the met-low or tumor-free groups in both models (Fig. 1A-B). Of note, in the breast carcinoma model, LSK differences did not reach statistical significance. In addition, their levels were substantially lower than those in the melanoma model, probably due to known technical difficulty of Sca1+ immunostaining in BALB/c mice (Spangrude and Brooks, 1993).

**Figure 1:**
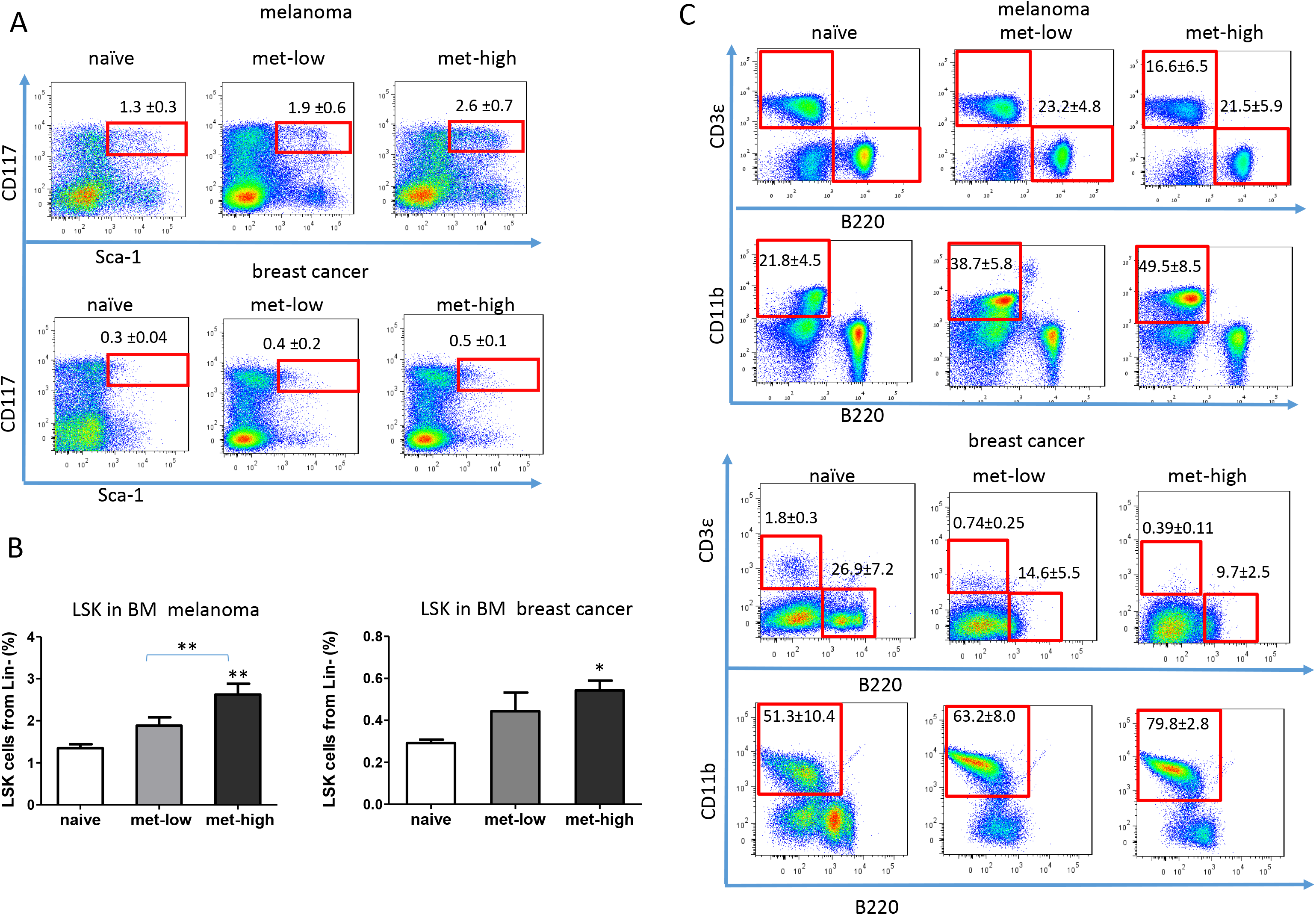
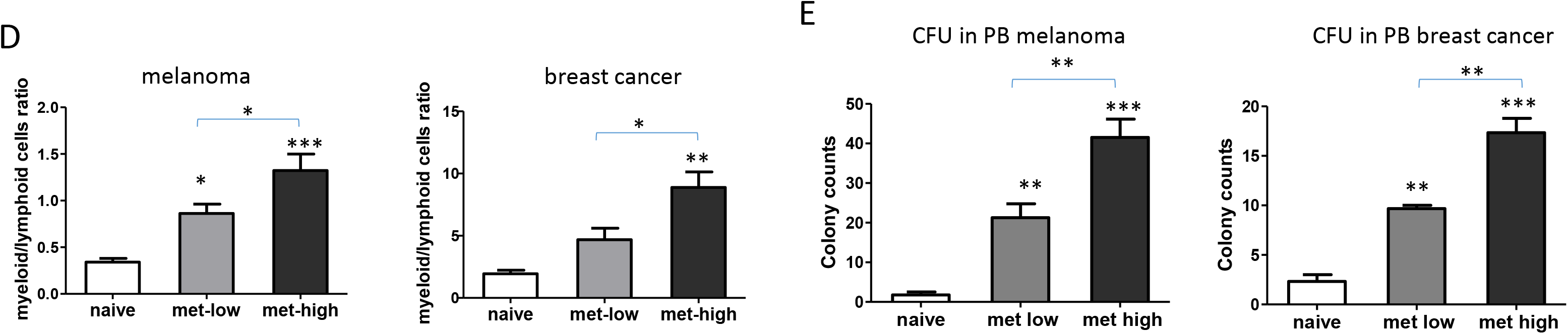
Metastatic tumors promote myeloid-biased skewing of HSPCs. Mice were either sub-dermally implanted in the flank with met-low B16-F1 or met-high B16-F10 melanoma cells (5×10^5^ cell/mouse), or implanted in the mammary fat pad with met-low 67NR or met-high 4T1 breast carcinoma cells (5×10^5^ cell/mouse). Control mice (naïve) were not implanted with tumor cells. At endpoint (day 18 and day 21, respectively), mice were sacrificed, and bone marrow and blood were harvested. (A) LSK cells in the BM were analyzed by flow cytometry. The cells were previously gated to exclude doublets and lineage positive cells. Representative flow cytometry plots are shown. (B) Quantification of LSK cells from the lineage negative cells (n=5-8 mice/group). (C) Lymphoid T and B cells and myeloid CD11b+ cells were quantified in the BM by flow cytometry. Representative plots are shown. Percentages are presented as means ± standard deviation (n=5-8 mice/group). (D) Myeloid to lymphoid cell ratios in the bone marrow compartment are shown (n=5-8 mice/group). (E) Circulating hematopoietic stem and progenitor cells in peripheral blood (PB) were evaluated by a colony forming assay. CFU colony counts are shown (n=3-4 mice/group). Statistical significance was assessed by one-way ANOVA, followed by Tukey post-hoc test when comparing more than two groups or unpaired two-tailed t-test when comparing two groups. Asterisks represent significance from control, unless indicated otherwise in the figure. Significant p values are shown as *, p<0.05, ** p<0.01, ***p<0.001.

Next, since tumor progression is associated with pronounced myelopoiesis (Wu et al., 2014), with the focus on the BM niche, we quantified myeloid and lymphoid cells in the BM compartment of naïve and tumor-bearing mice. Tumor-bearing mice exhibited an increase in myeloid cells and a decrease in lymphoid-derived B and T cells, in comparison to the tumor-free control group, in a comparable way to their changes in the tumor mass, as previously reported (Wildes et al., 2019). Importantly, the increased ratio of myeloid to lymphoid derived cells in the BM compartment was most pronounced in mice implanted with met-high tumors (Fig. 1C-D). Of note, mice implanted with met-high tumors exhibited the highest number of colony forming units (CFUs) derived from HSPCs in peripheral blood, further indicating increased hematopoiesis (Fig. 1E). Taken together, our findings suggest that tumors promote hematopoiesis, an effect which is augmented in highly metastatic tumors.

### Metastatic tumors dictate a long-lived education of HSPCs into myeloid-biased

We next asked whether tumor-induced myelopoiesis is due to myeloid-biased HSPC differentiation. To test this, LSK cells obtained from naïve GFP-expressing mice were mixed with either met-low or met-high melanoma tumor cells in Matrigel and subsequently implanted in recipient mice (Fig. 2A). Control recipient mice were implanted with Matrigel containing either LSK cells or tumor cells (met-low or met-high cells). After 10 days, tumor size and the differentiation of GFP+ cells in tumors, peripheral blood and bone marrow were assessed. Tumor size was significantly increased in mice implanted with the mixture of HSPCs and met-high tumor cells, an effect which was not statistically significant in the corresponding met-low tumor group (Fig. 2B). These results suggest that HSPCs give rise to hematopoietic cell subsets that support tumor growth. When comparing between met-high and met-low tumors in the context of GFP+ HSPC differentiation, no significant differences were detected in the percentages of GFP+ granulocytes, total macrophages and pro-inflammatory M1 macrophages. However, the level of tumor-supporting GFP+ M2 macrophages was significantly increased by approximately 2-fold in met-high tumors (Fig. 2C-F). No GFP+ cells were detected in peripheral blood or BM (data not shown). In the same experimental setting using Lin-cells instead of LSK cells, a similar trend was found in tumor growth and number of Lin- derived macrophages in mice bearing met-high tumors compared to met-low tumors. However, Lin- derived granulocytes exhibited increased numbers only in met-high tumors (Fig. S2A-D). Taken together, our results demonstrate that in met-high tumors, HSPCs undergo myeloid-biased, an effect known to promote tumor aggressiveness and metastasis.

**Figure 2:**
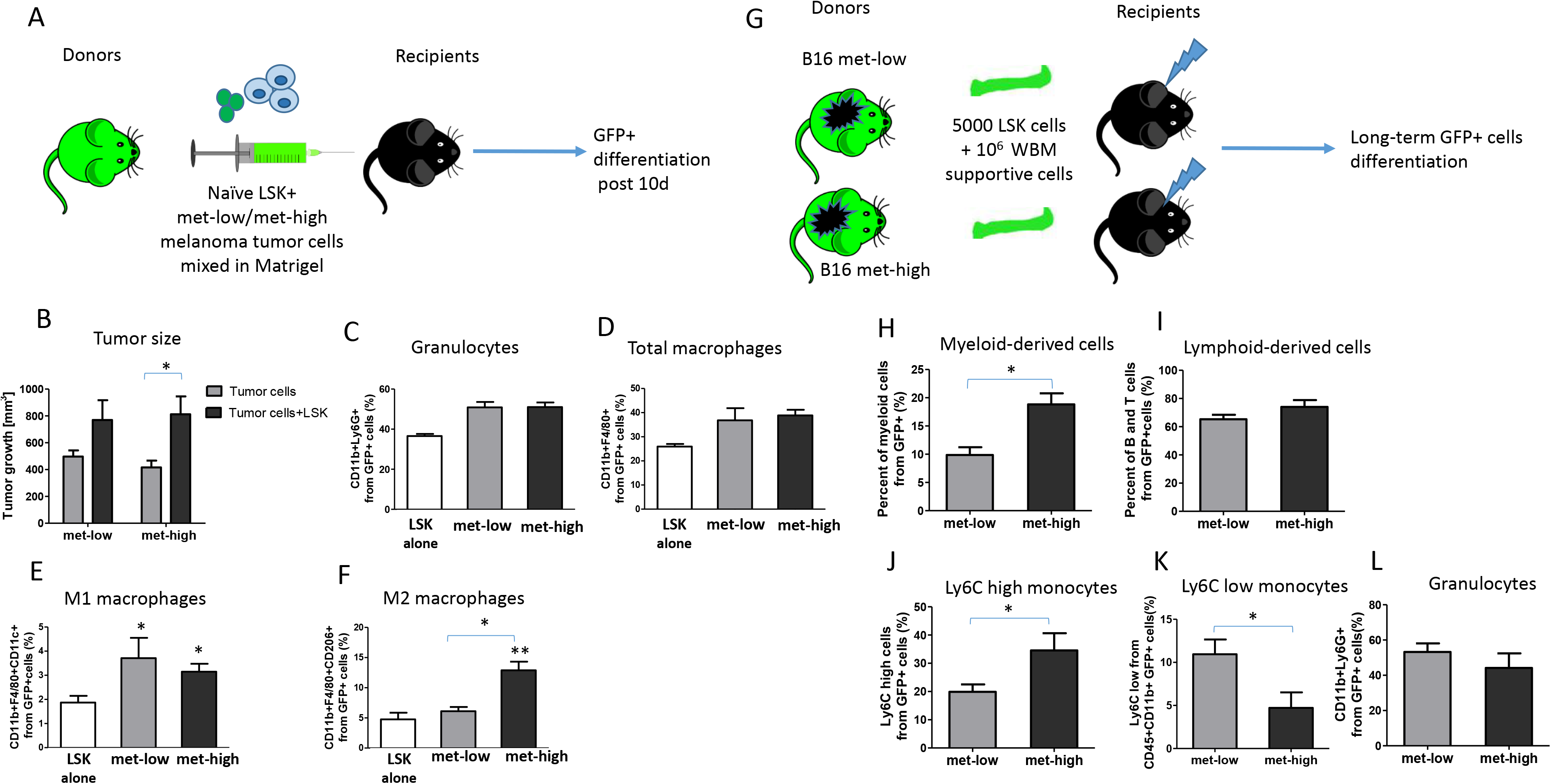
Metastatic tumors dictate HSPC differentiation. (A) A schematic representation of the LSK and tumor cell co-implantation experiment is shown. LSK cells obtained from naïve GFP-expressing donor mice were co-injected with either met-low or met-high melanoma tumor cells to recipient mice. Control recipient mice (not shown) were either injected with LSK cells alone, or with tumor cells alone. At endpoint, mice were sacrificed, and tumors harvested. GFP+ progeny in tumors was assessed by flow cytometry (n=4-5 mice/group). (B) Tumor volumes at endpoint are presented. (C-F) The percentages of granulocytes (C), total macrophages (D), M1 (E) and M2 (F) macrophages in tumors were determined by flow cytometry on GFP+ gated cells. (G) A schematic representation of the tumor-educated LSK transplantation experiment is shown. LSK cells were obtained from the bone marrow of donor mice harboring met-low or met-high melanomas. The LSK cells (5×10^3^) were injected together with whole bone marrow supportive cells (1×10^6^) from naïve mice into lethally-irradiated naïve recipient mice. Donor engraftment was monitored for 16 weeks post transplantation. GFP+ progeny in peripheral blood was assessed by flow cytometry. (H-I) Lineage distribution of myeloid (H) and lymphoid-derived cells (I) are shown. (J-L) The percentage of GFP+Ly6C^high^ monocytes (J), Ly6C^low^ monocytes (K) and GFP+Ly6G+ granulocytes (L) were gated from myeloid CD11b+ cells (n=5 mice/group). Statistical significance was assessed by one-way ANOVA, followed by Tukey post-hoc test when comparing more than two groups or unpaired two-tailed t-test when comparing two groups. Asterisks represent significance from control, unless indicated otherwise in the figure. Significant p values are shown as * p<0.05; ** p<0.01; ***p<0.001.

Next, to study the long-lived education and differentiation of HSPC by tumor cells, LSK or Lin-cells obtained from BM of GFP-expressing mice harboring either met-low or met-high melanoma tumors were transplanted along with whole BM (GFP-) supporting cells into lethally irradiated naive recipient mice (Fig. 2G and Fig. 2SE, respectively). After 16 weeks, the percentages of circulating GFP+ myeloid and lymphoid cells in the peripheral blood was evaluated. A significant increase in the percentage of myeloid but not lymphoid cells was found in mice transplanted with BM cells originating from met-high donors (Fig. 2H-I and Fig. S2F-G). Moreover, in the met-high group, the pro-inflammatory Ly6C^high^ monocyte fraction was significantly increased in tested GFP+ myeloid cells in peripheral blood, while there was a decrease in Ly6C^low^ monocytes that exhibit a low inflammatory profile (Fig. 2J-K). No difference was observed in the granulocyte population (Fig. 2L). These results indicate that met-high tumors dictate a long-lived direct differentiation of HSC towards myeloid cells, particularly to M2 macrophages in tumors and pro-inflammatory monocytes in peripheral blood.

To further strengthen the correlation between HSPC differentiation and immune cell composition in the tumor microenvironment of met-high and met-low tumors, we used high throughput mass cytometry (CyTOF) to analyze the percentage of different myeloid-derived cells in the stroma of met-high or met-low tumors followed by flow cytometry validation of specific cell types for both melanoma and breast carcinoma. Overall, the analysis of both CyTOF and flow cytometry data revealed a significant enrichment in M2 macrophages in met-high tumors compared to met-low tumors, whereas the levels of pro-inflammatory M1 macrophages were significantly decreased. In addition, the levels of Ly6C^high^ monocytes were increased in met-high tumors in comparison to met-low tumors. Moreover, the granulocyte population was also significantly increased in met-high tumors in the melanoma but not breast carcinoma model (Fig. S3). Overall, these results demonstrate similar trends to those found in the LSK bone marrow transplantation experiment, indicating that met-high tumors dictate a specific differentiation of HSPCs towards myeloid cells, especially M2 macrophages.

### Metastatic tumors affect myeloid progenitor composition towards MDPs

To further substantiate our hypothesis of HSPC programming by highly metastatic cells, we performed scRNA-seq of LSK cells obtained from the BM of mice implanted with met-low or met-high tumors. We next identified the specific cellular composition of the LSK cells by correlating single-cell expression profiles with ImmGen sorted cell reference transcriptomic database (Fig. 3A) (Aran et al., 2019; Heng et al., 2008). Specifically, we successfully identified myeloid progenitor cells including monocyte-dendritic and granulocyte-monocyte progenitor cells whose annotation were found to be significant (Fig. S4A, mean *P* = 0.0152 and 0.0079, respectively). Importantly, among the LSK differentiation pattern enriched cells, monocyte-dendritic progenitors (MDPs) were highly enriched in met-high tumor group whereas granulocyte monocyte progenitor (GMPs) were highly enriched in met-low tumor group (P < 0.0001, for both MDP and GMP, BCa bootstrap independent two-samples test for proportion differences) (Fig 3B-D). Furthermore, scRNA-seq data revealed that CX3CR1 is enriched within the MDP cells of the met-high tumor group (Fig. S4B-C *P* = 0.041, Wilcoxon rank-sum test), in line with a previous study demonstrating that MDPs are initially identified as CX3CR1+ cells (Yanez et al., 2017). Of note, when focusing on the tumor microenvironment, a significant elevation in the percentage of MDPs was observed in met-high melanoma and breast tumors compared to met-low tumors, while all other annotated cells including GMPs were not significantly changed (Fig. 3E and Fig. S4D-I).

**Figure 3:**
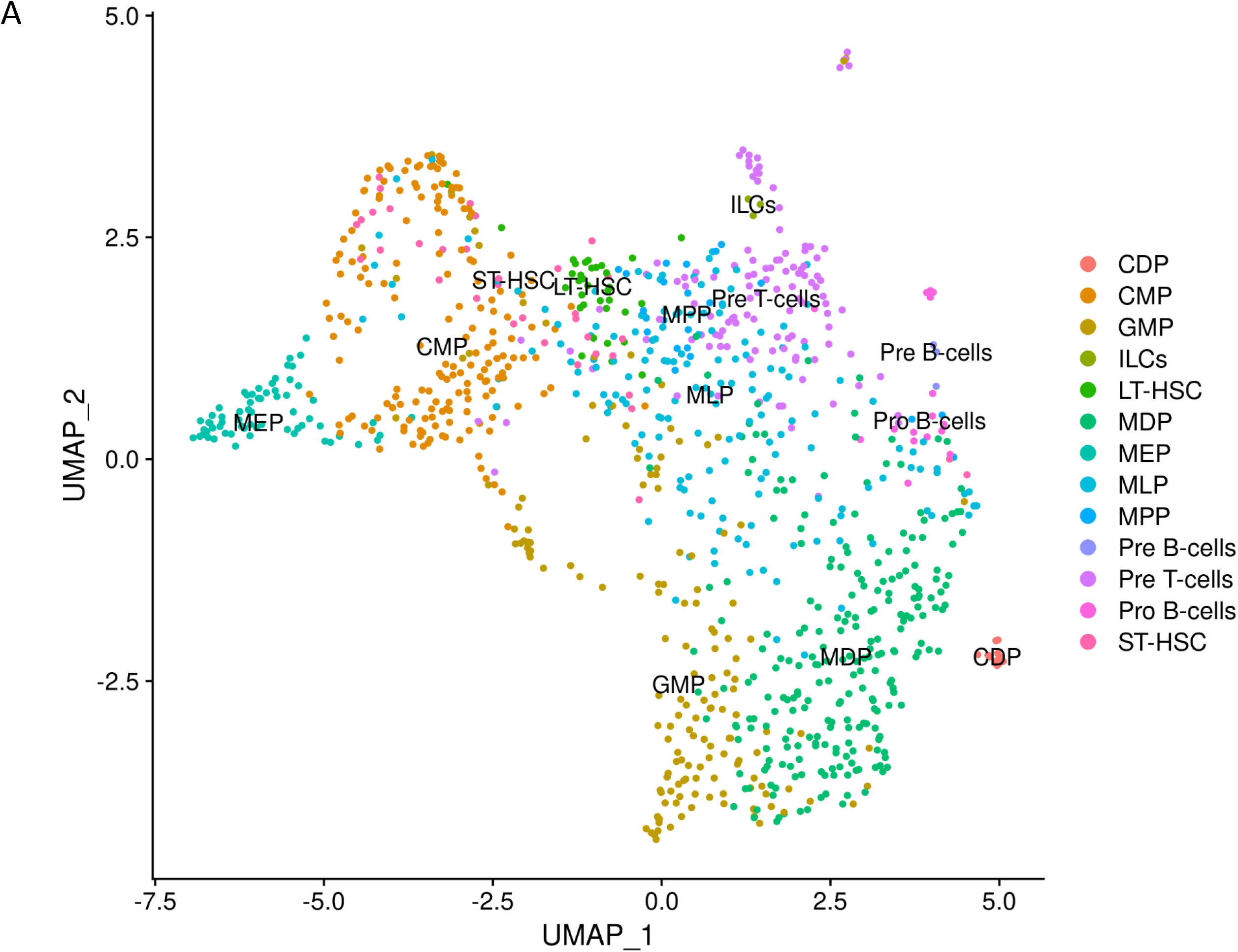
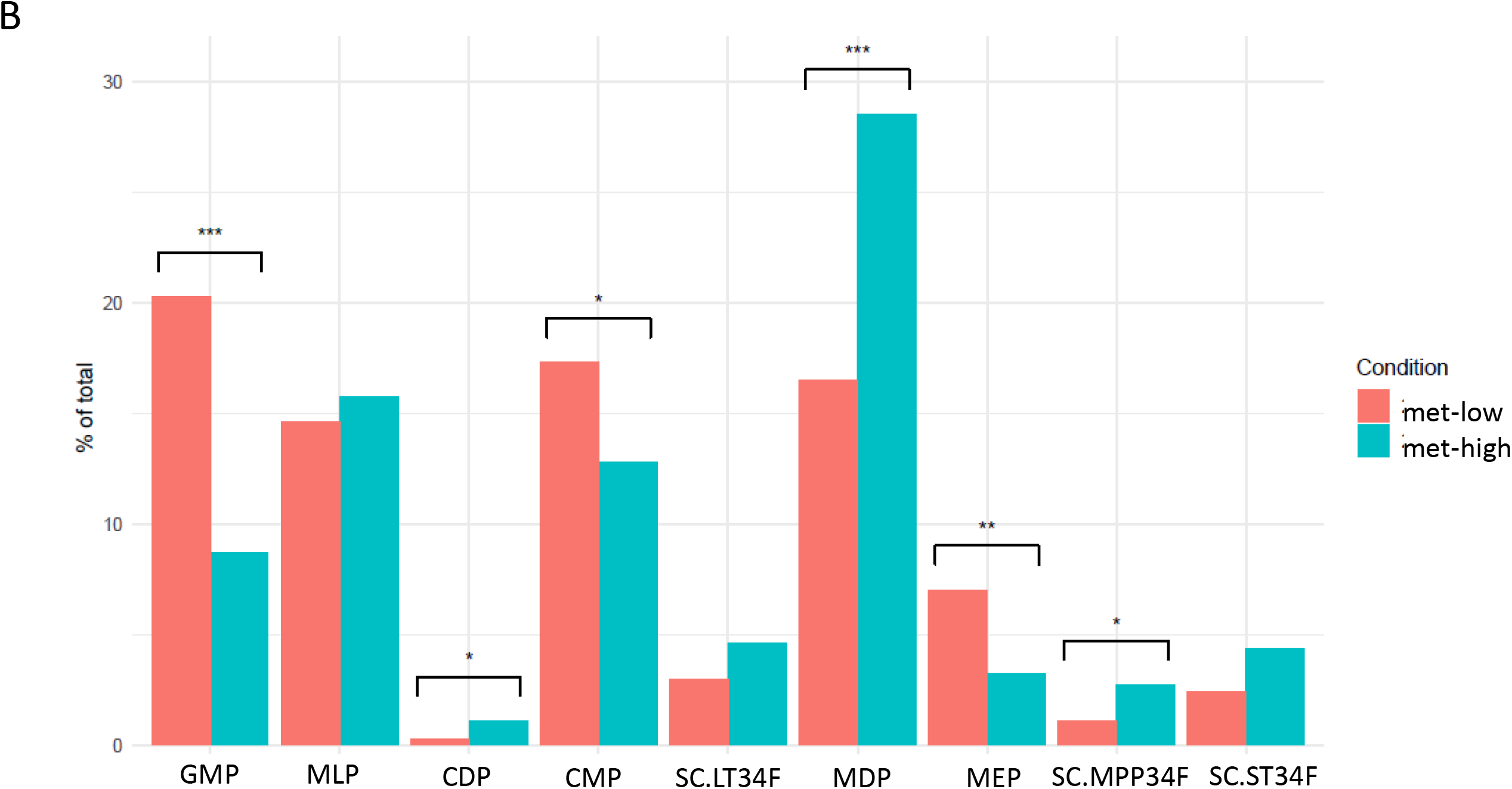
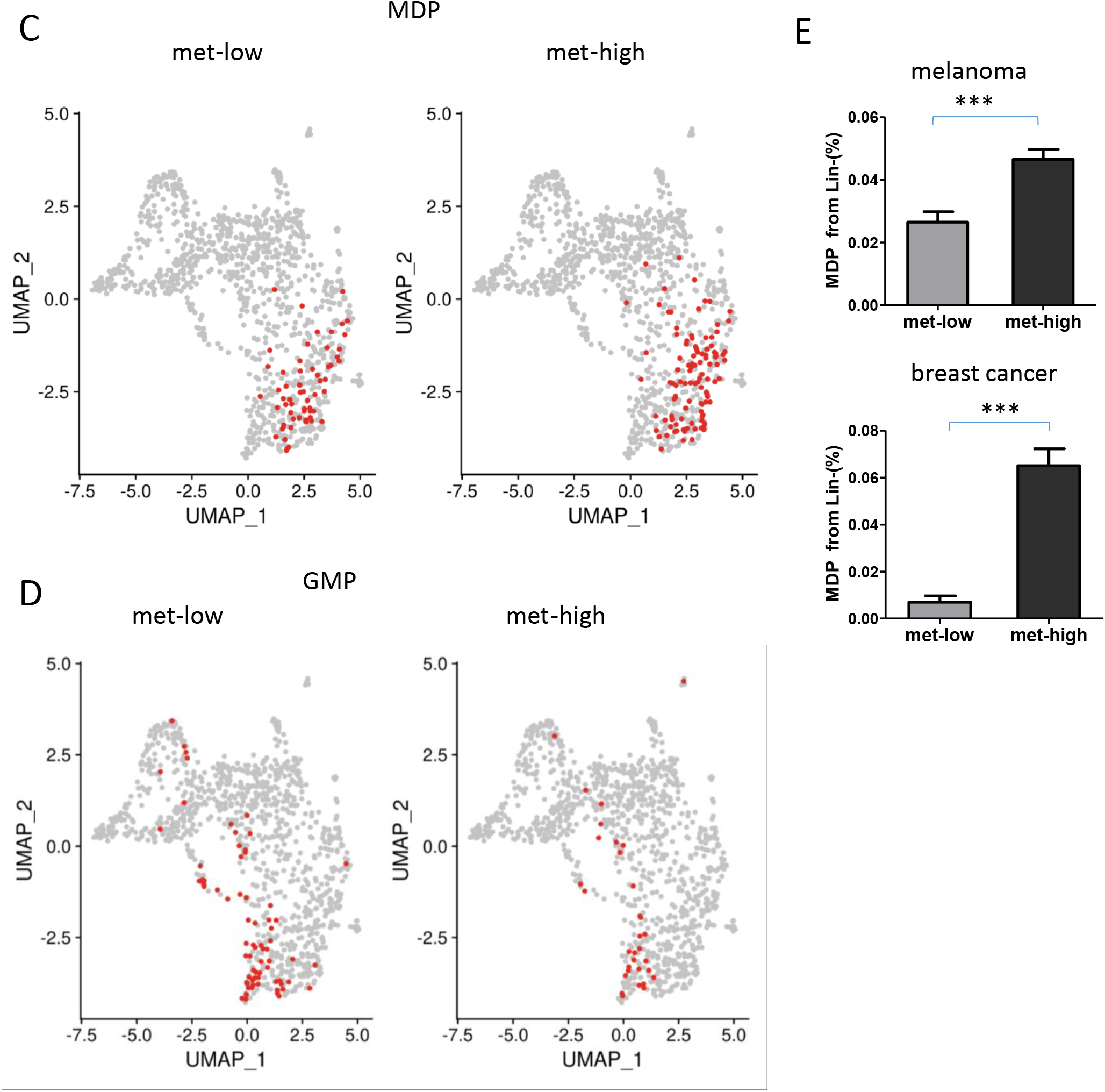
Metastatic tumors affect myeloid progenitor composition. Mice were implanted with met-high or met-low melanoma tumor cells. At endpoint, LSK cells were obtained from bone marrow and analyzed by single cell RNA sequencing. (A) UMAP plots of LSK clustering and individual cell type annotation by SingleR for both groups, using ImmGen gene expression compendium as a reference. MDP and GMP clusters are indicated by blue and red arrows, respectively. (B) Stem and progenitor enrichment levels in met-low and met-high groups, calculated by BCa bootstrap independent two-samples test for proportion differences. (C-D) Mapping of MDP (C) and GMP (D) cells on UMAP plots in met-low and met-high groups. Statistical significance was assessed by Wilcoxon rank-sum test. (E) Flow cytometry validation of MDP percentage from Lin-cells of met-low and met-high tumor bearing mice. Melanoma and breast carcinoma models are shown (n=5 mice/group in melanoma model, n=5-10 mice/group in breast carcinoma model). Statistical significance was assessed using unpaired two-tailed t-test and shown * p<0.05; ***p<0.001.

Next, to further study the functional differences of MDPs in met-high and met-low tumors, we analyzed differentially expressed genes (DEGs) in the MDP population in the BM of tumor-free, met-low, and met-high melanoma bearing mice. Genes differentially expressed in naïve and met-low MDPs (vs. met-high MDPs) significantly overlap (N=62, Fig. S4J). They are involved in similar immune-related pathways (Fig. S4K) and are potentially controlled by identical transcription factors (Fig. S4L). Notably, the serum response factor (SRF) network - known to be involved in myeloid-dendritic differentiation (Guenther et al., 2019), is enriched in naïve and met-low MDPs. Loss of SRF results in accumulation of undifferentiated HSCs (Ragu et al., 2010) - consistent with increased LSK levels within the bone marrow of met-high tumor bearing mice. In contrast, met-high MDPs are enriched for E2F-target genes involved in mitosis and the G2M checkpoint, suggesting they possess higher proliferative potential. Overall, these results suggest that met-low MDPs remain “naïve-like”, and that met-high tumors, specifically educate HSPCs by enriching the MDP population and altering its differentiation program.

### IL-6 is a key player in the differentiation of HSPCs into MDPs

We hypothesized that the crosstalk between tumor and bone marrow cells requires messenger proteins, e.g., cytokines. With the focus on MDPs, we searched for DEGs in MDPs between met-high and met-low groups. Overall, 207 genes were significantly altered between the two groups as shown in the volcano plot, including receptors and transmembrane proteins, such as IL-6Ra and IL-2R (Fig. S5A and Table S3). We next sought to identify tumor-secreted factors that correlate with the sc-RNA-seq data. A cytokine array to quantify the levels of a broad range of factors in the tumor-derived conditioned medium (TCM) of met-high and met-low melanomas was used. Several factors, including CXCL1, CXCL10, IL-3, IL-6 and IL-13, were found at higher levels in the TCM of met-high tumors (Fig. S5B). We then matched the cytokines to their corresponding receptors mentioned above. Using this approach, IL-6 and its receptor were found to be good candidates for mediating the crosstalk between tumor cells and HSPCs (Fig. 4A). Verification by ELISA revealed that IL-6 was significantly upregulated in met-high TCM in both melanoma and breast carcinoma models (Fig. 4B). The IL-6 receptor (IL-6Ra) was found to be upregulated in MDPs of mice harboring met-high melanoma (Fig. 4C; average fold change = 1.21, *P* = 0.02, Wilcoxon rank-sum test), and its expression is abundant in the MDP population shown in the UMAPplot (Fig. 4D). These results were verified by flow cytometry for both melanoma and breast carcinoma (Fig. S5C). Furthermore, to strengthen the involvement of IL-6 in met-high tumors, transcription signatures of CD45+ stroma cells obtained from tumor microenvironment, assessed by NanoString approach, revealed 7 upregulated genes, including THBS1, MAPKAPK2, CD9, TXNIP, CD24, ITGA5 and SYK (Fig. S5D). Importantly, some of these genes (THBS1, MAPKAPK2, CD9, TXNIP) were strongly associated with IL-6 signaling (Lim et al., 2020; Lopez-Dee et al., 2011; Shi et al., 2017), indicating that IL-6 or its associated genes are coupled with metastatic tumors, in alignment with our previous findings. Overall, these findings may explain the increased myeloid bias in met-high compared to met-low tumors.

**Figure 4:**
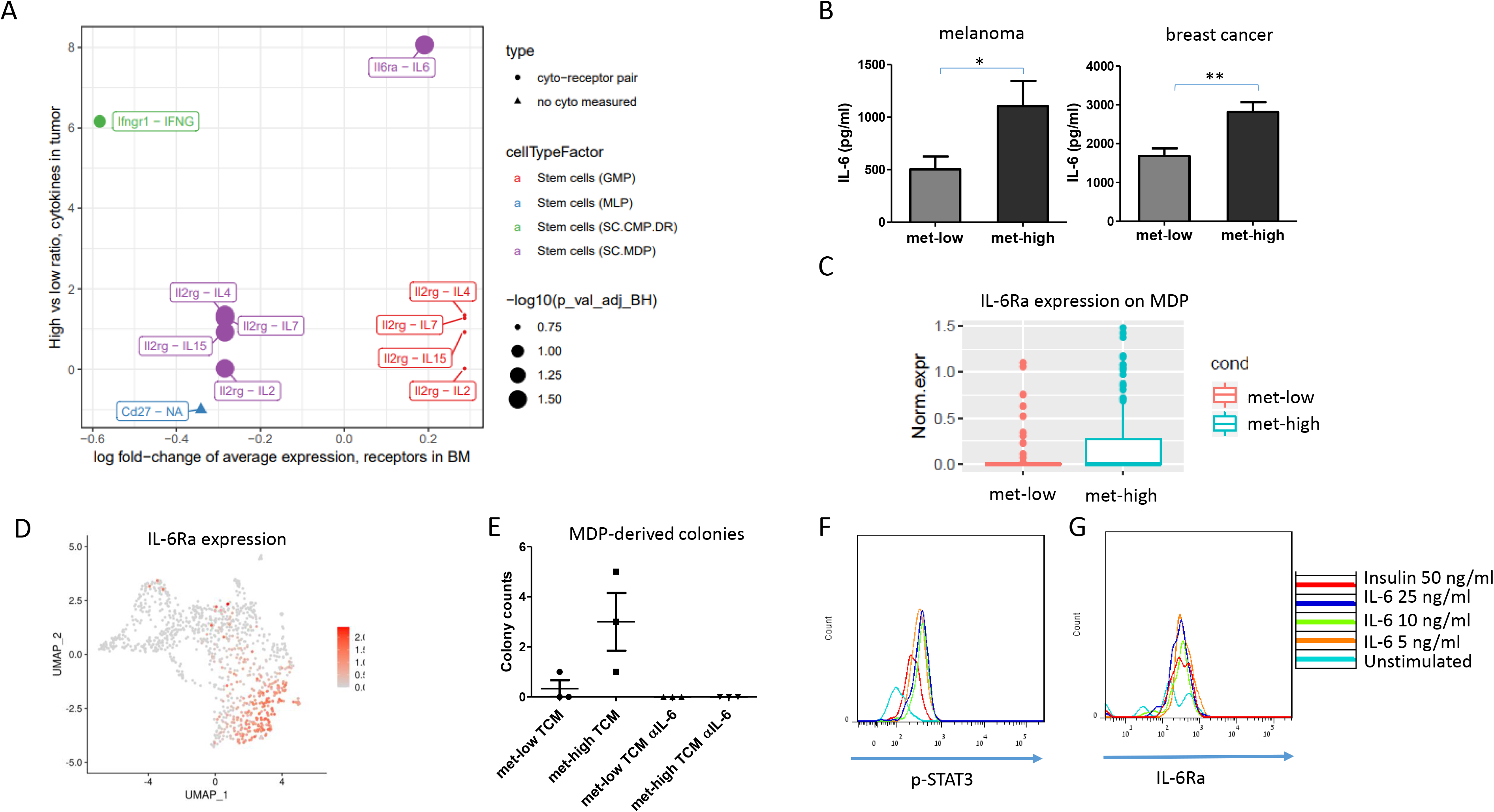
IL-6 mediates crosstalk between tumors and MDPs. (A) Mapping upregulated cytokines in melanoma met-high tumor conditioned medium (TCM) with their corresponding receptors, differentially expressed by progenitor populations, as identified by single cell RNA sequencing (Wilcoxon rank-sum test followed by Benjamini–Hochberg correction). (B) IL-6 levels were quantified by ELISA in met-low and met-high TCM of melanoma and breast carcinoma models (n= 5-8=mice / group). (C) IL-6Ra expression in the MDP population in the bone marrow of met-low and met-high melanoma bearing mice, based on scRNA-seq dataset (average fold change = 1.21, *P* = 0.02, Wilcoxon rank-sum test). (D) Single cell IL-6Ra mRNA expression labeling on the UMAP plot. (F) MDPs from naïve mice were grown in Methocult medium supplemented with met-low or met-high melanoma TCM in the presence or absence of anti-IL-6 neutralizing antibodies. MDP-derived colonies were counted (n=3 biological repeats / group). (E-F) MDPs obtained from the bone marrow of naïve mice were stimulated with escalating doses of IL-6. The levels of p-STAT3 (E) and IL-6Ra (F) were determined by flow cytometry. Statistical significance was assessed by one-way ANOVA, followed by Tukey post-hoc test when comparing more than two groups or unpaired two-tailed t-test when comparing two groups. Significant p values are shown as * p<0.05; ** p<0.01.

To directly assess the effect of IL-6 on MDP proliferation and differentiation, MDP cells isolated from BM of naïve mice were seeded in Methocult medium supplemented with met-high or met-low TCM in the presence or absence of neutralizing IL-6 antibody. Following 12 days of incubation, myeloid colonies were scored. MDP cells formed more monocyte colonies in the presence of met-high TCM in comparison to the met-low group. Importantly, neutralizing IL-6 in the met-high group completely abolished MDP growth and colony formation (Fig. 4E).

One of the downstream effects in the IL-6 pathway is phosphorylation of STAT3 (Heinrich et al., 2003). To verify that MDPs respond directly to IL-6, we stimulated BM cells obtained from naïve mice with escalating doses of IL-6 for 30 minutes and evaluated p-STAT3 expression specifically in the MDP population. Indeed, p-STAT3 levels in MDP cells were increased upon stimulation with IL-6 (Fig. 4F), demonstrating that IL-6 directly affects IL-6Ra-expressing-MDPs (Fig. 4G). Taken together, our results suggest that IL-6 secreted by highly metastatic tumors directly promotes myeloid-biased HSPC differentiation into MDPs.

### IL-6 promotes metastasis via MDP-derived macrophages

We next sought to investigate the effect of IL-6 signaling in vivo focusing on primary tumor growth, metastasis and HSPC differentiation. To this end, mice were implanted with met-low or met-high melanoma or breast carcinoma cells, and subsequently either treated with IgG control or anti-IL-6 antibodies. The rates of primary tumor growth were similar in all groups throughout the course of the experiment (Fig. S6A-B). However, in both tumor models, anti-IL-6 treatment dramatically reduced the incidence of lung metastasis in mice harboring met-high tumors (Fig. 5A-D). This effect was accompanied by a significant decrease in MDP levels in the BM of these mice (Fig. 5E-F), suggesting a correlation between MDP levels and IL-6-mediated metastasis.

**Figure 5:**
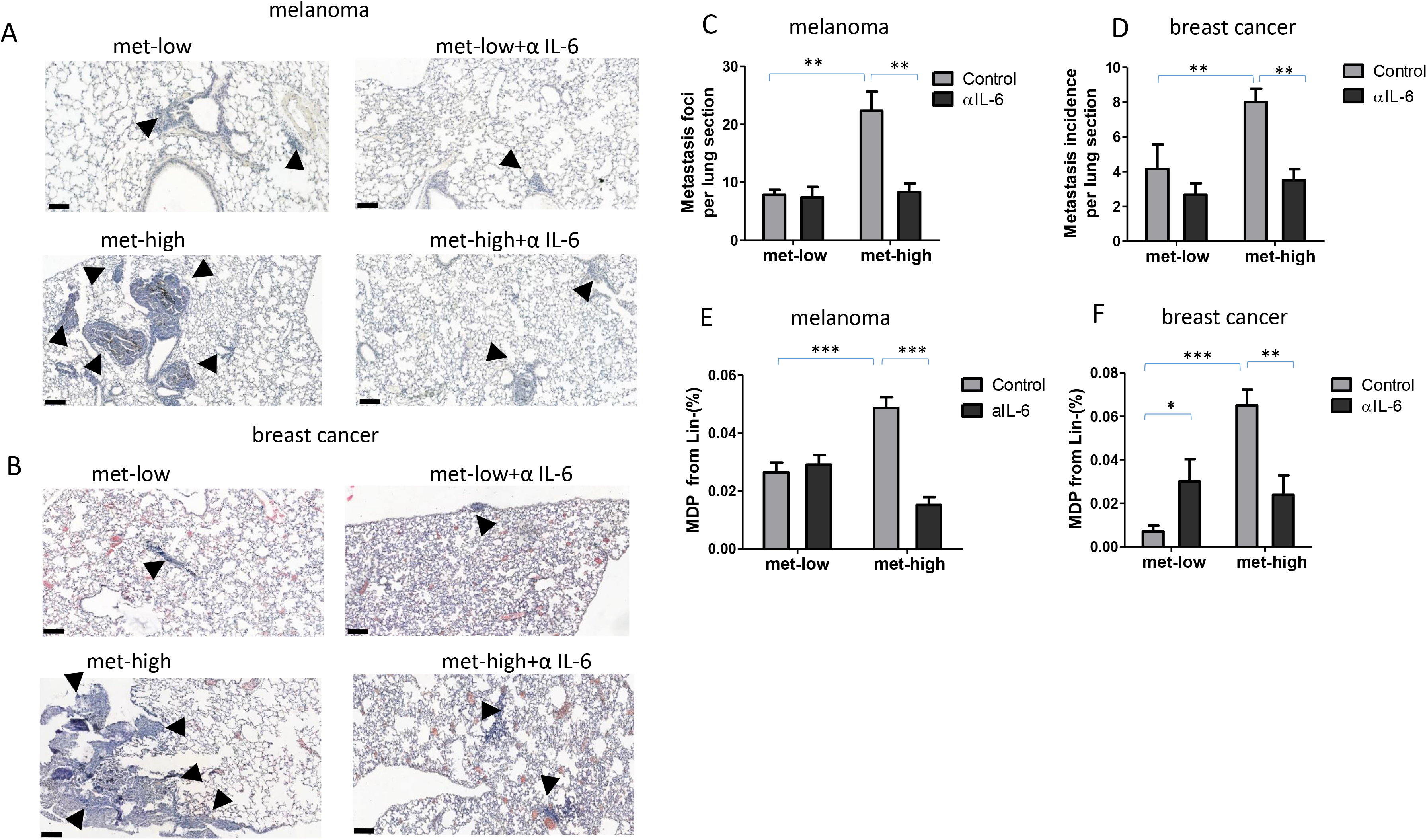
IL-6 pathway blockade in MDPs inhibits metastasis. Mice were implanted with met-low or met-high melanoma or breast carcinoma cells. One week later, mice were treated with IgG (control) or anti-IL-6 antibodies twice weekly. At endpoint, mice were sacrificed, lungs were removed, and bone marrow was harvested (n=5-8 mice/group). (A-B) Representative images of lung sections are shown, bar=100μm. Arrows indicate metastatic foci. (C-D) Metastatic foci per lung section were quantified (n=5-7 sections/mouse). (E-F) MDP levels in BM of melanoma met-low and met-high melanoma (E) or breast carcinoma (F) were assessed by flow cytometry. Statistical significance was assessed by one-way ANOVA, followed by Tukey post-hoc test when comparing more than two groups. Significant p values are shown as * p<0.05; ** p<0.005.

We next directly tested the effect of tumor-induced education of MDPs on metastasis in vivo by performing an MDP adoptive transfer experiment. To this end, GFP-expressing donor mice were implanted with met-high melanoma cells and then treated with control IgG or anti-IL-6 antibodies. GFP+ MDPs from these mice were then adoptively transferred into recipient mice subsequently injected with met-low tumor cells, as schematically illustrated in Fig. 6A. No significant difference in tumor growth was observed between the two groups (Fig. 6B). The number of metastatic lesions in the lungs of mice adoptively transferred with MDPs obtained from anti-IL-6-treated mice was significantly reduced in comparison to MDP from IgG-treated treated control mice (Fig. 6C-D). However, when GMP cells that can differentiate to monocytes (Yanez et al., 2017), were adoptively transferred to mice bearing met-low tumors, as illustrated in Fig. S6C, no significant difference in lung metastasis were observed (Fig. S6D-E), suggesting that MDPs but not GMPs contribute to metastasis.

**Figure 6:**
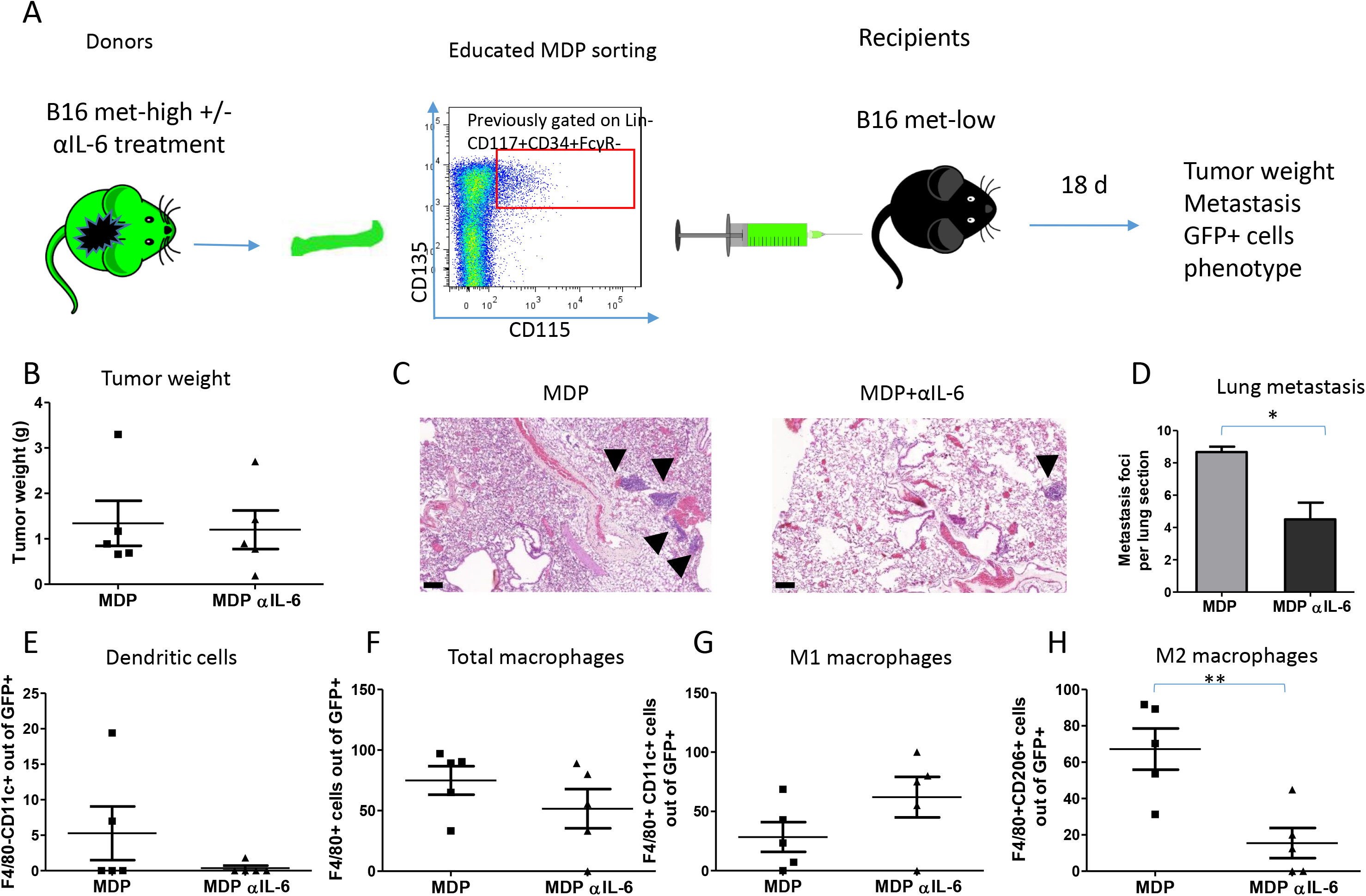
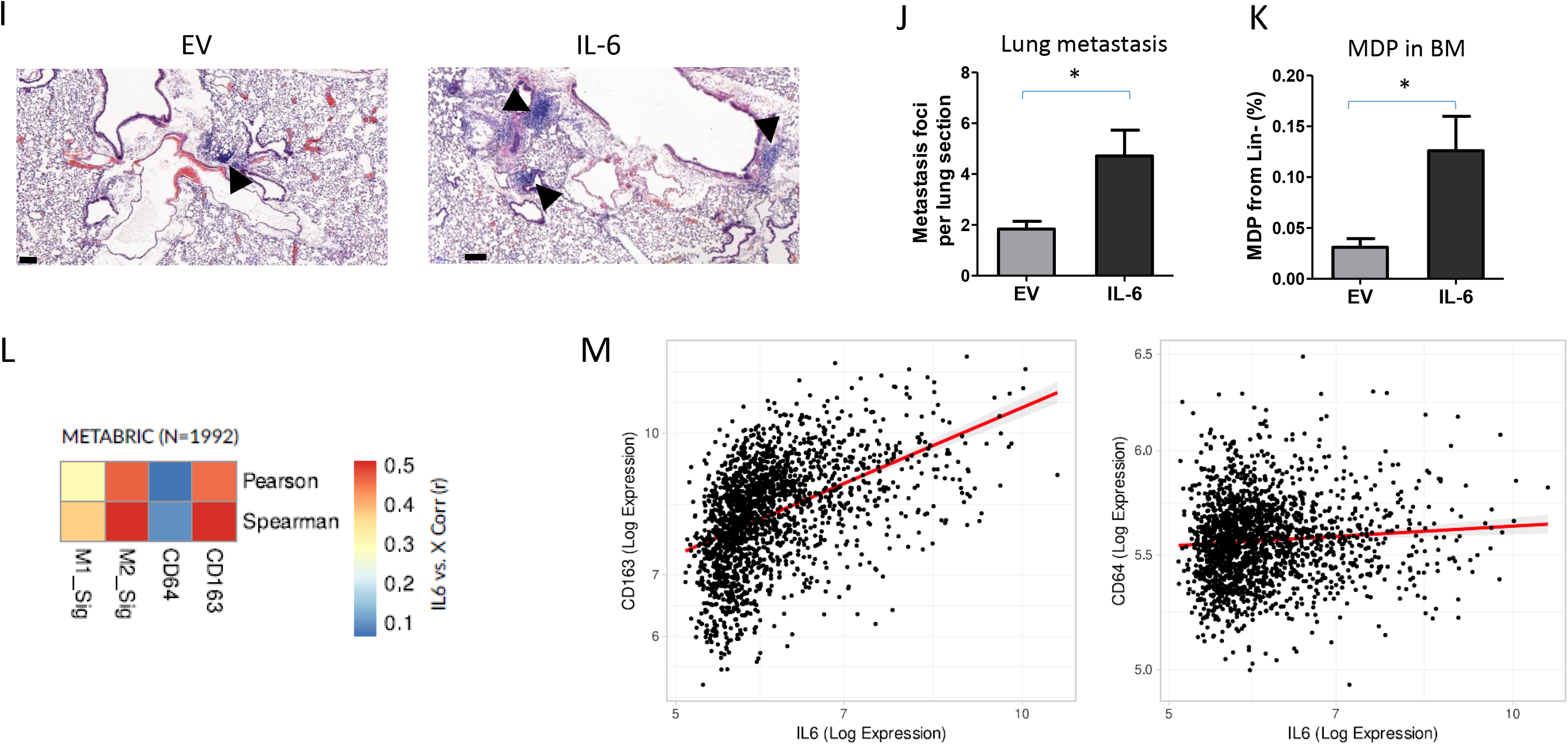
IL-6 promotes metastasis via MDP-derived macrophages. (A) A schematic representation of the experimental design is shown. GFP-expressing mice implanted with met-high melanoma cells were either treated with IgG control or anti-IL-6 antibodies twice weekly. At endpoint, FACS sorted MDPs were intravenously injected into naïve recipient mice (n=5 mice / group). The next day, the recipient mice were implanted with met-low melanoma cells. At endpoint, mice were sacrificed, and tumors and lungs were extracted. (B) Tumor weights are shown. (C) Representative images of lung sections are shown, bar=100μm. Arrows indicate metastatic foci. (D) Metastatic foci per lung section were quantified (n=5-7 sections per group). (E-H) GFP+ cells in single cell suspensions of lung samples were evaluated by flow cytometry to quantify the percentage of dendritic cells (E), total macrophages (F), M1 macrophages (G), and M2 macrophages (H). (I-K) Eight-to-ten week old C57Bl/6 mice were implanted with B16 IL-6 overexpressing cells or with corresponding control EV cells. At endpoint (day 18), mice were sacrificed, lungs were removed, and the bone marrow was harvested (n=4-6 mice/group). (I) Representative images of lung sections are shown, bar=100μm. Arrows indicate metastatic foci. (J) Metastatic foci per lung section were quantified (n=4-6 sections/mouse). (K) MDP levels in BM were assessed by flow cytometry. (L) Heat-map of linear correlations in r values between IL-6 expression and M1 or M2 gene signatures as well as single genes CD64 or CD163 representing M1 and M2 subsets, respectively. The data was obtained from human breast cancer samples using the METABRIC dataset. (M) Scatter graphs of IL-6 expression in correlation with CD64 or CD163 in human breast cancer samples (METABRIC dataset). Statistical significance was assessed by unpaired two-tailed t-test. Significant p values are shown as * p<0.05; ** p<0.01.

Since MDPs give rise to monocytes and dendritic cells (Morales et al., 1995), we next evaluated these two populations in lung single cell suspensions. Dendritic cells were barely detected in the GFP+ MDP cell populations (Fig 6E). There were no differences in the levels of GFP+ total macrophages and M1 macrophages between the two groups (Fig. 6F-G). However, the percentage of M2 GFP+ macrophages was significantly decreased in the lungs of mice adoptively transferred with MDPs obtained from anti-IL-6-treated mice, indicating that IL-6 blockade induces a functional change in MDPs (Fig. 6H). We should note that naïve GMP population, cultured in Methocult, supplemented with met-low or met-high TCM in the presence or absence of ant-IL-6, revealed low growth of colonies (Fig. S6F). Overall, these findings suggest that MDPs rather that GMPs, upon IL-6 secretion from highly metastatic tumors, differentiate into M2 macrophages at the metastatic sites and promote metastasis.

### IL-6-induced MDP differentiation is a driver protein of the metastatic switch

To demonstrate that IL-6 is responsible for a metastatic switch in our experimental settings, we generated met-low melanoma tumors overexpressing IL-6 or empty vector (EV) control (B16-F1-IL-6 and B16-F1-EV, respectively). IL-6 expression in cultured cells was verified by ELISA (Fig. S7A), and the two cell lines exhibited similar proliferation rates (Fig. S7B). Next, mice were implanted with B16-F1-IL-6 or B16-F1-EV and tumor growth was assessed. There was no significant difference in tumor growth between the two groups (Fig. S7C), while a significant increase in IL-6 was found in the TCM of B16-F1-IL-6 group compared to B16-F1-EV (Fig. S7D), suggesting that IL-6 has no direct pro-tumorigenic role in the primary tumor microenvironment. However, the incidence of lung metastasis was significantly increased in B16-F1-IL-6 tumor bearing mice (Fig. 6I-J). Moreover, the elevation in lung metastasis was directly associated with increased MDP levels detected in the bone marrow (Fig. 6K), while the levels of GMPs, CMPs, and MEPs remained unchanged between the two groups (Fig. S7E-G). These results further indicate that IL-6 contributes to the metastatic switch mediated in part by HSPC differentiation into MDPs that further differentiate into M2-macrophage supporting metastasis.

To further correlate between MDP and macrophages supporting metastasis in human samples, we analyzed RNA-seq from breast carcinoma patients using the METABRIC dataset (Curtis et al., 2012). A moderate, positive correlation (r=~0.4-0.5) was found between IL-6 expression and the molecular signature of M2 macrophages (CD163, CD204, CD206, CD200R1, TGM2, IL1R2), whereas little correlation was found with an M1 gene signature (CD64, MHCII, CD86, CD80, CD68, NOS2) (Fig. 6L). This trend is strengthened when considering a single marker gene largely unique to each macrophage subtype (Fig. 6M), suggesting that a similar mechanism may exist in humans. Overall, these findings suggest that IL-6 directly affects MDPs, which in turn, differentiate into metastasis-promoting cells, an effect which may also present in humans.

## Discussion

The hematopoietic system, in addition to its role in replenishing the blood cell compartment, also contributes to the immune response in various pathological conditions. Tumor progression is linked to pronounced perturbations in myelopoiesis, similar to inflammation, resulting in increased levels of committed and early myeloid progenitor populations (Wu et al., 2014){Hoggatt, 2016 #8583}{Wilson, 2008 #8590}. During tumor-induced myelopoiesis, HSPCs migrate from the BM and accumulate at distant sites. There, they differentiate into tumor-supporting myeloid cells to drive immunosurveillance (Gabrilovich et al., 2012), participate in the formation of the pre-metastatic niche (Giles et al., 2016; Kaplan et al., 2005), and facilitate tumor cell seeding (Kaplan et al., 2005). A recent study has demonstrated that early progenitors can be found in the blood circulation of patients with different types of solid tumors, with higher levels of GMPs that eventually differentiate into granulocytes. Moreover, increased levels of myeloid progenitors correlated with overall poor outcome in these patients(Wu et al., 2014). In agreement, we observed myeloid bias in peripheral blood associated with highly metastatic tumors, as demonstrated by BM transplantation experiments using educated Lin- and LSK cells obtained from met-low or met-high tumor bearing mice. Specifically, scRNA-sequencing of LSK cells obtained from mice harboring met-high tumors revealed a gene signature that reflects an MDP differentiation pattern. These cells were then found to serve as the origin of tumor-associated macrophages in the pulmonary metastatic sites. These findings strongly support the concept that met-high tumors affect the HSPC progeny to support metastatic-promoting accessory cells. Importantly, these effects were absent in met-low tumors which resulted in gene signature that is closely related to naïve HSPCs. Moreover, the bone marrow transplantation experiment using educated LSK cells from tumor bearing mice further suggests that HSPC fate is long-lived as they maintain their biased progeny in bone marrow of reconstituted mice over a three month period. In fact, sustained met-high educated HSPC specific differentiation suggests the involvement of epigenetic modifications that might regulate their programming, a process that should be further elucidated. Furthermore, while there is an overwhelming amount of evidence for HSPCs giving rise to MDSCs in tumor conditions (Wildes et al., 2019), we found that LSK cells educated by met-low or met-high tumor conditions exhibit an MDSC phenotype, with specific differentiation program which further gives rise to M2 macrophages only in met-high tumors. Indeed, studies have demonstrated that myeloid cells and tumor-associated immunosuppressive macrophages (M2 macrophages) home to the pre-metastatic sites where they contribute to the recruitment and retention of circulating tumor cells (Condamine et al., 2015; Doak et al., 2018; Giles et al., 2016). In agreement with these studies, here we provide further evidence for the origin of M2 macrophages, since we demonstrate that highly metastatic tumors promote metastasis through enrichment of MDP-derived macrophages localized at the metastatic sites. We further demonstrate that these effects exist only in met-high tumors but not in met-low tumors, suggesting a direct involvement of tumor cells on HSPC progeny. The existence of two distinct and independently regulated cellular pathways challenge the function of GMP-derived vs. MDP-derived monocytes and their relative contribution to metastasis. Here we suggest that MDPs but not GFPs have a novel function in metastasis progression.

HSPCs express receptors of various inflammatory cytokines, such as IL-1, IL-6, interferon ⍰ and TLR ligands, making them sensitive to external stimuli (King and Goodell, 2011; Nagai et al., 2006; Rieger et al., 2009). It has been suggested that such receptors regulate HSPC differentiation. Thus, tumors may use this opportunity to condition the BM compartment via factors secreted by the tumor microenvironment. Our experimental model revealed an association between upregulated IL-6 levels in met-high TCM and elevated expression levels of IL-6Ra in met-high educated MDPs. The IL-6/IL-6Ra pathway is often hyperactivated in many types of cancer (Johnson et al., 2018) and plays a key role in proliferation, survival and invasiveness of tumor cells (Chang et al., 2013; Yu et al., 2009). In addition, it has been shown that IL-6 mediates metastatic mechanisms such as stimulation of epithelial-to-mesenchymal transition (Rokavec et al., 2014), and induction of matrix metalloproteinases (MMPs) (Yu and Jove, 2004). Our study provides another role for IL-6 in metastasis. We show that IL-6 induces metastasis through programming HSPCs towards tumor-supporting metastatic cells, with no effect on the primary tumor. Importantly, we demonstrate that IL-6 education of MDPs alone enhances the metastatic potential of tumor cells. Specifically, IL-6-educated MDPs adoptively transferred into met-low tumor bearing mice, resulted in increased metastasis, in a similar phenotype found in met-high tumors. These results may hold in human, as we revealed a moderate correlation between IL-6 and gene signature of M2 but not M1 macrophages in human breast cancer samples from the METABRIC dataset. Thus, IL-6 should be further tested as a prognostic biomarker for metastasis in the context of immune cell composition in tumors and metastatic sites.

In the present study, anti-IL-6 treatment resulted in a dramatic drop in lung metastasis in met-high tumor bearing mice, while no effect was observed in the met-low group. We showed that anti-IL-6 treatment reduced the levels of MDPs in the BM along with M2 macrophages in the lungs of met-high tumor bearing mice, suggesting that MDP-derived macrophages serve as key promoters of metastasis formation. Anti-IL-6 monoclonal therapy was recently approved by the FDA for Castelman disease patients (Johnson et al., 2018). In the context of solid tumors, despite success in pre-clinical studies (Johnson et al., 2018), the efficacy of such treatment was limited in Phase I-II clinical trials among patients with prostate, ovarian and lung cancers (Angevin et al., 2014; Karkera et al., 2011). Our results further support these clinical studies, since we demonstrated that anti-IL-6 treatment does not inhibit primary tumor growth, whereas it dramatically reduces metastatic burden. In the clinic, anti-IL-6 studies were performed in patients with advanced metastatic disease. Our study therefore suggests that evaluating plasma levels of IL-6 at early stages, before metastases appear, probably in the neoadjuvant setting, may provide better insights into the therapeutic potential of IL-6 inhibition.

In summary, our study demonstrates a novel mechanism underlying direct tumor-mediated dictation of HSPC differentiation and programming of its progeny in the tumor microenvironment. This, in turn, becomes an essential process for tumor progression and metastasis. We demonstrate that the crosstalk between the tumor and distant BM niche is mediated by IL-6/IL-6Ra signaling, paving the way towards new therapeutic approaches to target metastasis.

## Conflict of interest

The authors declare no conflict of interest.

## Author contribution

Conception and design: KMK, KK, SSS and YS.

Acquisition of data: KMK, KK, TC, RN, MT, ZR, and BJ.

Analysis and interpretation of data: KMK, KK, TC, RN, TZ, RG, ZAR, SSS, and YS.

Writing, review, and/or revision of the manuscript: KMK, KK, TC, RG, ZAR, SSS, and YS.

Study supervision: YS.

## Acknowledgment

This work is supported primarily by H2020 European Research Council Grant (771112) given to YS. TC is supported by TICC-Rubinstein fellowship. This study makes use of data generated by the Molecular Taxonomy Breast Cancer International Consortium. Funding for the project was provided by Cancer Research UK and the British Columbia Cancer Agency Branch.

## Notes

### Competing Interest Statement

The authors have declared no competing interest.

### Summary of Updates

The revised manuscript fix errors related to the definition of the cells, e.g., MDPs and hematopoietic stem and progenitor cells (HSPCs).

## References

Amir el, A.D., K.L. Davis, M.D. Tadmor, E.F. Simonds, J.H. Levine, S.C. Bendall, D.K. Shenfeld, S. Krishnaswamy, G.P. Nolan, and D. Pe’er. 2013. viSNE enables visualization of high dimensional single-cell data and reveals phenotypic heterogeneity of leukemia. Nat Biotechnol 31:545–552.

Angevin, E., J. Tabernero, E. Elez, S.J. Cohen, R. Bahleda, J.L. van Laethem, C. Ottensmeier, J.A. Lopez-Martin, S. Clive, F. Joly, I. Ray-Coquard, L. Dirix, J.P. Machiels, N. Steven, M. Reddy, B. Hall, T.A. Puchalski, R. Bandekar, H. van de Velde, B. Tromp, J. Vermeulen, and R. Kurzrock. 2014. A phase I/II, multiple-dose, dose-escalation study of siltuximab, an anti-interleukin-6 monoclonal antibody, in patients with advanced solid tumors. Clinical cancer research: an official journal of the American Association for Cancer Research 20:2192–2204.

Aran, D., A.P. Looney, L. Liu, E. Wu, V. Fong, A. Hsu, S. Chak, R.P. Naikawadi, P.J. Wolters, A.R. Abate, A.J. Butte, and M. Bhattacharya. 2019. Reference-based analysis of lung single-cell sequencing reveals a transitional profibrotic macrophage. Nat Immunol 20:163–172.

Butler, A., P. Hoffman, P. Smibert, E. Papalexi, and R. Satija. 2018. Integrating single-cell transcriptomic data across different conditions, technologies, and species. Nat Biotechnol 36:411–420.

Chang, Q., E. Bournazou, P. Sansone, M. Berishaj, S.P. Gao, L. Daly, J. Wels, T. Theilen, S. Granitto, X. Zhang, J. Cotari, M.L. Alpaugh, E. de Stanchina, K. Manova, M. Li, M. Bonafe, C. Ceccarelli, M. Taffurelli, D. Santini, G. Altan-Bonnet, R. Kaplan, L. Norton, N. Nishimoto, D. Huszar, D. Lyden, and J. Bromberg. 2013. The IL-6/JAK/Stat3 feed-forward loop drives tumorigenesis and metastasis. Neoplasia 15:848–862.

Chang, Y.H., T.Y. Chu, and D.C. Ding. 2020. Human fallopian tube epithelial cells exhibit stemness features, self-renewal capacity, and Wnt-related organoid formation. J Biomed Sci 27:32.

Condamine, T., I. Ramachandran, J.I. Youn, and D.I. Gabrilovich. 2015. Regulation of tumor metastasis by myeloid-derived suppressor cells. Annu Rev Med 66:97–110.

Curtis, C., S.P. Shah, S.F. Chin, G. Turashvili, O.M. Rueda, M.J. Dunning, D. Speed, A.G. Lynch, S. Samarajiwa, Y. Yuan, S. Graf, G. Ha, G. Haffari, A. Bashashati, R. Russell, S. McKinney, A. Langerod, A. Green, E. Provenzano, G. Wishart, S. Pinder, P. Watson, F. Markowetz, L. Murphy, I. Ellis, A. Purushotham, A.L. Borresen-Dale, J.D. Brenton, S. Tavare, C. Caldas, and S. Aparicio. 2012. The genomic and transcriptomic architecture of 2,000 breast tumours reveals novel subgroups. Nature 486:346–352.

Doak, G.R., K.L. Schwertfeger, and D.K. Wood. 2018. Distant Relations: Macrophage Functions in the Metastatic Niche. Trends Cancer 4:445–459.

Finak, G., A. McDavid, M. Yajima, J. Deng, V. Gersuk, A.K. Shalek, C.K. Slichter, H.W. Miller, M.J. McElrath, M. Prlic, P.S. Linsley, and R. Gottardo. 2015. MAST: a flexible statistical framework for assessing transcriptional changes and characterizing heterogeneity in single-cell RNA sequencing data. Genome Biol 16:278.

Gabrilovich, D.I. 2017. Myeloid-Derived Suppressor Cells. Cancer immunology research 5:3–8.

Gabrilovich, D.I., S. Ostrand-Rosenberg, and V. Bronte. 2012. Coordinated regulation of myeloid cells by tumours. Nat Rev Immunol 12:253–268.

Giles, A.J., C.M. Reid, J.D. Evans, M. Murgai, Y. Vicioso, S.L. Highfill, M. Kasai, L. Vahdat, C.L. Mackall, D. Lyden, L. Wexler, and R.N. Kaplan. 2016. Activation of Hematopoietic Stem/Progenitor Cells Promotes Immunosuppression Within the Pre-metastatic Niche. Cancer Res 76:1335–1347.

Guenther, C., I. Faisal, L.M. Uotila, M.L. Asens, H. Harjunpaa, T. Savinko, T. Ohman, S. Yao, M. Moser, S.W. Morris, S. Tojkander, and S.C. Fagerholm. 2019. A beta2-Integrin/MRTF-A/SRF Pathway Regulates Dendritic Cell Gene Expression, Adhesion, and Traction Force Generation. Front Immunol 10:1138.

Heinrich, P.C., I. Behrmann, S. Haan, H.M. Hermanns, G. Muller-Newen, and F. Schaper. 2003. Principles of interleukin (IL)-6-type cytokine signalling and its regulation. Biochem J 374:1–20.

Heng, T.S., M.W. Painter, and C. Immunological Genome Project. 2008. The Immunological Genome Project: networks of gene expression in immune cells. Nat Immunol 9:1091–1094.

Hoggatt, J., Y. Kfoury, and D.T. Scadden. 2016. Hematopoietic Stem Cell Niche in Health and Disease. Annu Rev Pathol 11:555–581.

Ishiguro, T., M. Nakajima, M. Naito, T. Muto, and T. Tsuruo. 1996. Identification of genes differentially expressed in B16 murine melanoma sublines with different metastatic potentials. Cancer Res 56:875–879.

Johnson, D.E., R.A. O’Keefe, and J.R. Grandis. 2018. Targeting the IL-6/JAK/STAT3 signalling axis in cancer. Nat Rev Clin Oncol 15:234–248.

Kanehisa, M., Y. Sato, M. Kawashima, M. Furumichi, and M. Tanabe. 2016. KEGG as a reference resource for gene and protein annotation. Nucleic Acids Res 44:D457–462.

Kaplan, R.N., R.D. Riba, S. Zacharoulis, A.H. Bramley, L. Vincent, C. Costa, D.D. MacDonald, D.K. Jin, K. Shido, S.A. Kerns, Z. Zhu, D. Hicklin, Y. Wu, J.L. Port, N. Altorki, E.R. Port, D. Ruggero, S.V. Shmelkov, K.K. Jensen, S. Rafii, and D. Lyden. 2005. VEGFR1-positive haematopoietic bone marrow progenitors initiate the pre-metastatic niche. Nature 438:820–827.

Karkera, J., H. Steiner, W. Li, V. Skradski, P.L. Moser, S. Riethdorf, M. Reddy, T. Puchalski, K. Safer, U. Prabhakar, K. Pantel, M. Qi, and Z. Culig. 2011. The anti-interleukin-6 antibody siltuximab down-regulates genes implicated in tumorigenesis in prostate cancer patients from a phase I study. Prostate 71:1455–1465.

King, K.Y., and M.A. Goodell. 2011. Inflammatory modulation of HSCs: viewing the HSC as a foundation for the immune response. Nat Rev Immunol 11:685–692.

Liberzon, A., C. Birger, H. Thorvaldsdottir, M. Ghandi, J.P. Mesirov, and P. Tamayo. 2015. The Molecular Signatures Database (MSigDB) hallmark gene set collection. Cell Syst 1:417–425.

Lim, J.O., N.R. Shin, Y.S. Seo, H.H. Nam, J.W. Ko, T.Y. Jung, S.J. Lee, H.J. Kim, Y.K. Cho, J.C. Kim, I.C. Lee, J.S. Kim, and I.S. Shin. 2020. Silibinin Attenuates Silica Dioxide Nanoparticles-Induced Inflammation by Suppressing TXNIP/MAPKs/AP-1 Signaling. Cells 9:

Lopez-Dee, Z., K. Pidcock, and L.S. Gutierrez. 2011. Thrombospondin-1: multiple paths to inflammation. Mediators Inflamm 2011:296069.

Love, M.I., W. Huber, and S. Anders. 2014. Moderated estimation of fold change and dispersion for RNA-seq data with DESeq2. Genome Biol 15:550.

Mehlen, P., and A. Puisieux. 2006. Metastasis: a question of life or death. Nature reviews. Cancer 6:449–458.

Morales, D.E., K.A. McGowan, D.S. Grant, S. Maheshwari, D. Bhartiya, M.C. Cid, H.K. Kleinman, and H.W. Schnaper. 1995. Estrogen promotes angiogenic activity in human umbilical vein endothelial cells in vitro and in a murine model. Circulation 91:755–763.

Nagai, Y., K.P. Garrett, S. Ohta, U. Bahrun, T. Kouro, S. Akira, K. Takatsu, and P.W. Kincade. 2006. Toll-like receptors on hematopoietic progenitor cells stimulate innate immune system replenishment. Immunity 24:801–812.

Ragu, C., G. Elain, E. Mylonas, C. Ottolenghi, N. Cagnard, D. Daegelen, E. Passegue, W. Vainchenker, O.A. Bernard, and V. Penard-Lacronique. 2010. The transcription factor Srf regulates hematopoietic stem cell adhesion. Blood 116:4464–4473.

Rieger, M.A., P.S. Hoppe, B.M. Smejkal, A.C. Eitelhuber, and T. Schroeder. 2009. Hematopoietic cytokines can instruct lineage choice. Science 325:217–218.

Rokavec, M., M.G. Oner, H. Li, R. Jackstadt, L. Jiang, D. Lodygin, M. Kaller, D. Horst, P.K. Ziegler, S. Schwitalla, J. Slotta-Huspenina, F.G. Bader, F.R. Greten, and H. Hermeking. 2014. IL-6R/STAT3/miR-34a feedback loop promotes EMT-mediated colorectal cancer invasion and metastasis. J Clin Invest 124:1853–1867.

Schmaus, A., and J.P. Sleeman. 2015. Hyaluronidase-1 expression promotes lung metastasis in syngeneic mouse tumor models without affecting accumulation of small hyaluronan oligosaccharides in tumor interstitial fluid. Glycobiology 25:258–268.

Shaked, Y., E. Pham, S. Hariharan, K. Magidey, O. Beyar-Katz, P. Xu, S. Man, F.T. Wu, V. Miller, D. Andrews, and R.S. Kerbel. 2016. Evidence Implicating Immunological Host Effects in the Efficacy of Metronomic Low-Dose Chemotherapy. Cancer Res 76:5983–5993.

Shi, Y., W. Zhou, L. Cheng, C. Chen, Z. Huang, X. Fang, Q. Wu, Z. He, S. Xu, J.D. Lathia, Y. Ping, J.N. Rich, X.W. Bian, and S. Bao. 2017. Tetraspanin CD9 stabilizes gp130 by preventing its ubiquitin-dependent lysosomal degradation to promote STAT3 activation in glioma stem cells. Cell Death Differ 24:167–180.

Spangrude, G.J., and D.M. Brooks. 1993. Mouse strain variability in the expression of the hematopoietic stem cell antigen Ly-6A/E by bone marrow cells. Blood 82:3327–3332.

van de Rijn, M., S. Heimfeld, G.J. Spangrude, and I.L. Weissman. 1989. Mouse hematopoietic stem-cell antigen Sca-1 is a member of the Ly-6 antigen family. Proc Natl Acad Sci U S A 86:4634–4638.

Wang, H., C. Horbinski, H. Wu, Y. Liu, S. Sheng, J. Liu, H. Weiss, A.J. Stromberg, and C. Wang. 2016. NanoStringDiff: a novel statistical method for differential expression analysis based on NanoString nCounter data. Nucleic Acids Res 44:e151.

Wang, Y., Y. Ding, N. Guo, and S. Wang. 2019. MDSCs: Key Criminals of Tumor Pre-metastatic Niche Formation. Front Immunol 10:172.

Wildes, T.J., C.T. Flores, and D.A. Mitchell. 2019. Concise Review: Modulating Cancer Immunity with Hematopoietic Stem and Progenitor Cells. Stem cells 37:166–175.

Wu, W.C., H.W. Sun, H.T. Chen, J. Liang, X.J. Yu, C. Wu, Z. Wang, and L. Zheng. 2014. Circulating hematopoietic stem and progenitor cells are myeloid-biased in cancer patients. Proc Natl Acad Sci U S A 111:4221–4226.

Yanez, A., S.G. Coetzee, A. Olsson, D.E. Muench, B.P. Berman, D.J. Hazelett, N. Salomonis, H.L. Grimes, and H.S. Goodridge. 2017. Granulocyte-Monocyte Progenitors and Monocyte-Dendritic Cell Progenitors Independently Produce Functionally Distinct Monocytes. Immunity 47:890–902 e894.

Yu, G., L.G. Wang, Y. Han, and Q.Y. He. 2012. clusterProfiler: an R package for comparing biological themes among gene clusters. OMICS 16:284–287.

Yu, H., and R. Jove. 2004. The STATs of cancer--new molecular targets come of age. Nature reviews. Cancer 4:97–105.

Yu, H., D. Pardoll, and R. Jove. 2009. STATs in cancer inflammation and immunity: a leading role for STAT3. Nature reviews. Cancer 9:798–809.

