## Supplemental online materials for "Tumor-educated monocyte-dendritic progenitors promote a metastatic switch"

### Supplemental Materials Online

Magidey-Klein et. al.,

### Supplementary methods

#### IL-6 overexpressing cells

Cells genetically modified to express IL-6 For overexpression of IL-6, B16-F1 cells were transfected with 1 µg DNA of pCMV3 vector encoding IL-6 or the empty control pCMV3 vector (EV) using PolyJet (SignaGen Laboratories, MD, USA) following the manufacturer's instructions. For generating stable clones, transfected cells were grown for two weeks with hygromycin (250µg/ml). Conditioned medium obtained after 24 hours was analyzed for IL-6 levels using ELISA. Cell proliferation of IL-6 overexpressing cells was carried out by XTT, as previously described(Chang et al., 2020).

Kerenyi, M.A. 2014. LT-HSC Methylcellulose Assay. *Bio Protoc* 4:

Liberzon, A., C. Birger, H. Thorvaldsdottir, M. Ghandi, J.P. Mesirov, and P. Tamayo. 2015. The Molecular Signatures Database (MSigDB) hallmark gene set collection. *Cell Syst* 1:417-425.

Love, M.I., W. Huber, and S. Anders. 2014. Moderated estimation of fold change and dispersion for RNA-seq data with DESeq2. *Genome Biol* 15:550.

Shaked, Y., E. Pham, S. Hariharan, K. Magidey, O. Beyar-Katz, P. Xu, S. Man, F.T. Wu, V. Miller, D. Andrews, and R.S. Kerbel. 2016. Evidence Implicating Immunological Host Effects in the Efficacy of Metronomic Low-Dose Chemotherapy. *Cancer Res* 76:5983-5993.

Wang, H., C. Horbinski, H. Wu, Y. Liu, S. Sheng, J. Liu, H. Weiss, A.J. Stromberg, and C. Wang. 2016. NanoStringDiff: a novel statistical method for differential expression analysis based on NanoString nCounter data. *Nucleic Acids Res* 44:e151.

Yu, G., L.G. Wang, Y. Han, and Q.Y. He. 2012. clusterProfiler: an R package for comparing biological themes among gene clusters. *OMICS* 16:284-287.

### Supplemental tables and figures online

**Supplemental Table S1**

| Population | Markers |
|---|---|
| LSK | Lineage-Sca-1+CD117+ |
| GMP | Lineage-Sca-1-CD117+CD34-FcγR+ |
| CMP | Lineage-Sca-1-CD117+CD34+FcγR- |
| MEP | Lineage-Sca-1-CD117+CD34- FcγR- |
| MDP | Lineage-Sca-1-CD117+CD34+FcγR-CD115+FLT3+ |
| Granulocytes | CD11b+Ly6G+/Gr-1+ |
| Macrophages | Cd11b+F4/80+Ly6G- |
| M1 macrophages | Cd11b+F4/80+CD11c+Ly6G- |
| M2 macrophages | CD11b+ F4/80+CD206+Ly6G- |
| Inflammatory monocytes | CD11b+Ly6C$^{high}$Ly6G$^{low}$ CD115+ |
| Anti-inflammatory monocytes | CD11b+Ly6C$^{low}$Ly6G- |
| Dendritic cells | Cd11b+F4/80+Ly6G-MHCII+ |
| B cells | CD45+B220+ |
| T cells | CD45+CD3ε+ |

**Supplemental Table S2**

| Metal | Ab |
|---|---|
| 141Pr | CD80 |
| 142Nd | GR1 |
| 143Nd | CD86 |
| 144Nd | F4/80 |
| 145Nd | CD4 |
| 146Nd | CD45R |
| 147Sm | Ly6c |
| 148Nd | CD138 |
| 149Sm | CD8 |
| 150Nd | Ly6g |
| 151Eu | CD48 |
| 152Sm | CD90 |
| 153Eu | CD14 |
| 154Sm | CD11c |
| 155Gd | CD45 |
| 156Gd | CD49b |
| 157Gd | CD19 |
| 158Gd | CD34 |
| 159Tb | CD27 |
| 160Gd | CD69 |
| 161Dy | CD150 |
| 162Dy | TCRb |
| 163Dy | CD127 |
| 164Dy | CD28 |
| 165Ho | CD115 |
| 166Er | CD133 |
| 167Er | CD93 |
| 168Er | CD117 |
| 169Tm | CD79b |
| 170Er | CD62L |
| 171Yb | CD44 |
| 172Yb | CD43 |
| 173Yb | Sca-1 |
| 174Yb | Vegfr2 |
| 175Lu | CD5 |
| 176Yb | Cd11b |

**Figure S1. Rates of tumor growth and metastasis in met-high and met-low melanoma and breast tumor models.** C57BL/6 mice were sub-dermally implanted in the flank with met-low B16-F1 or met-high B16-F10 melanoma cells. BALB/c mice were implanted in the mammary fat pad with met-low 67NR or met-high 4T1 breast carcinoma cells (n=5 mice/group). Tumor growth was monitored regularly. At endpoint, lungs were removed and assessed for metastasis. (A) Tumor growth curves are presented. (B) Representative lung sections are shown, bar=100μm. Arrows indicate metastatic foci. (C) Metastatic foci per lung section were quantified (n=5 sections/mouse). Significant p values are shown as ** p<0.01, as assessed by unpaired two-tailed t-test.

**Figure S2. Lineage negative cells contribute to tumor growth and myelopoiesis.** (A) A schematic representation of the Lin- and tumor cell co-implantation experiment is shown. Lin- cells obtained from naïve GFP-expressing donor mice were co-injected in Matrigel with either met-low or met-high melanoma tumor cells to recipient mice. Control recipient mice (not shown) were either injected in Matrigel with Lin- cells alone, or with tumor cells alone. Tumor growth was monitored. At endpoint, mice were sacrificed, and tumors harvested. GFP+ progeny in tumors was assessed by flow cytometry (n=4-5 mice/group). (B) Tumor growth curves are shown. (C-D) The percentages of granulocytes (C) and total macrophages (D) in tumors were determined by flow cytometry on GFP+ gated cells. (E) A schematic representation of the tumor-educated Lin- transplantation experiment is shown. Lin- cells were obtained from the bone marrow of donor mice harboring met-low or met-high melanomas. The Lin- cells ($1x10^5$) were injected together with whole bone marrow supportive cells ($1x10^6$) from naïve mice into lethally-irradiated naïve recipient mice. Donor engraftment was monitored for 16 weeks post transplantation. GFP+ progeny in peripheral blood was assessed by flow cytometry. (F-G) The percentages of donor myeloid-derived (F) and lymphoid-derived cells (G) were assessed in peripheral blood. Statistical significance was assessed by one-way ANOVA, followed by Tukey post-hoc test when comparing more than two groups or unpaired two-tailed t-test when comparing two groups. Asterisks represent significance from control, unless indicated otherwise in the figure. Significant p values are shown as * p<0.05; ** p<0.01; *** p<0.001.

**Figure S3. Immune cell composition in met-high and met-low tumors.** (A-C) Mice were implanted with met-high or met-low tumor cells of melanoma (A-B) or breast carcinoma (C) models. At endpoint, tumors were removed and prepared as single cell suspensions. (A) viSNE plot obtained by immune cell CyTOF analysis is based on CD11b+ myeloid cells of met-low and met-high melanoma tumors (3 samples per group were pooled). (B-C) The percentages of macrophages and their M1 and M2 subsets, Ly6C$^{high}$, Ly6C$^{low}$ monocytes and granulocytes were analyzed by flow cytometry (n=5-8 mice for each group). Statistical significance was assessed by unpaired two-tailed t-test. Asterisks represent significance from control, unless indicated otherwise in the figure. Significant p values are shown as * p<0.05; ** p<0.01; *** p<0.001.

**Figure S4. LSK clustering in met-high and met-low tumors.** Mice were implanted with met-high or met-low melanoma tumor cells. At endpoint, LSK cells were obtained from bone marrow and analyzed by single cell RNA sequencing**.** (A) Cell type annotation significance levels (chi-squared outlier test for the top-scored cell type match). MDPs and GMPs are indicated by blue and red arrows, respectively. Statistical significance was assessed by Wilcoxon rank-sum test. (B-C) Single cell CX3CR1 mRNA expression labeling on the UMAP plot (B) and its expression specifically in MDP populations of met-low and met-high tumors (C). Statistical significance was assessed by Wilcoxon rank-sum test. (D-I) Flow cytometry validation of GMP (D,J,), CMP (E, ,H), and MEP (F, Ipercentages from Lin- cells of melanoma (D-F) or breast carcinoma (G-I) met-low and met-high tumor bearing mice (n=5 mice/group in melanoma model, n=5-10 mice/group in breast carcinoma model). Results are not statistically significant. (J) Venn diagram of differentially expressed genes (DEGs) in naïve and met-low MDPs vs met-high MDPs. (K) GSEA analysis of upregulated and downregulated biological processes comparing met-low MDPs or naïve MDPs vs met-high MDPs. (L) Transcription factors enrichment in met-low/naïve MDPs comparing with met-low MDPs.

**Figure S5. The secretion profile of met-high tumors is associated with IL-6.** (A) Volcano plot of significantly decreased (left side) and increased (right side) genes in MDP cells (met-high vs. met-low groups; melanoma model; Wilcoxon rank-sum test followed by Benjamini–Hochberg correction). (B) A cytokine array was performed to determine the levels of a range of factors in tumor conditioned medium (TCM) of met-high and met-low melanomas. Fold change is shown (met-high vs. met-low). (C) Representative plots of IL-6Ra expression on MDPs, obtained from bone marrow of met-low and met-high bearing mice, was assessed by flow cytometry. (D) Heat map of differentially expressed genes of CD45+ live cells, obtained from met-low and met-high melanoma tumors. Statistical significance was assessed by unpaired two-tailed t-test. Significant p values are shown as * p<0.05; ** p<0.01; *** p<0.001.

**Figure S6. GMP-derived cells do not support metastasis.** (A-B) Mice were implanted with met-low or met-high melanoma (A) or breast carcinoma (B) cells. One week later, mice were treated with IgG (control) or anti-IL-6 antibodies twice weekly (indicated by arrows). Tumor growth curves are shown (n=5-8 mice/group). (C) A schematic representation of the experimental design is shown. GFP-expressing mice were implanted with met-high melanoma cells. At endpoint, MDPs and GMPs were sorted by FACS, and were subsequently intravenously injected into naïve recipient mice (n=3-4 mice / group). The next day, the recipient mice were implanted with met-low melanoma cells. At end point, mice were sacrificed, and lungs were removed for the evaluation of metastasis. (D) Representative images of lung sections are shown, bar=100µm. Arrows indicate metastatic foci. (E) Metastatic foci per lung section were quantified (n=3-4 sections/mouse). (F) GMPs from naïve mice were grown in MethoCult medium supplemented with met-low or met-high melanoma tumor conditioned medium (TCM) in the presence or absence of anti-IL-6 neutralizing antibodies. GMP-derived colonies were counted (n=3 biological repeats / group). Statistical significance was assessed by one-way ANOVA, followed by Tukey post-hoc test when comparing more than two groups or unpaired two-tailed t-test when comparing two groups. Asterisks represent significance from control, unless indicated otherwise in the figure. Significant p values are shown as ** p<0.01.

**Figure S7. IL-6 overexpression in met-low tumors and its effect on HSPC lineage.** (A) Conditioned medium collected from IL-6 overexpressing and EV control B16-F1 cells was tested for IL-6 levels by ELISA. (B) Proliferation rate of IL-6 overexpressing cells and EV control cells was tested by XTT assay. (C-G) Eight-to-ten week old C57Bl/6 mice (n=4-6 mice/group) were implanted with B16-F1-IL-6 overexpressing cells or with corresponding control EV cells. Tumor growth was assessed (C). At endpoint (day 18), mice were sacrificed, tumors were removed and the bone marrow was harvested IL-6 levels were quantified in B16-F1 overexpressing IL-6 and B16-F1 EV control TCM, using ELISA (D). The levels of GMP (E), CMP (F), and MEP (G) from Lin- cells were assessed in the bone marrow of mice from both groups by flow cytometry. Statistical significance was assessed by unpaired two-tailed t-test. Significant p values are shown as * p<0.05.

Fig. S1

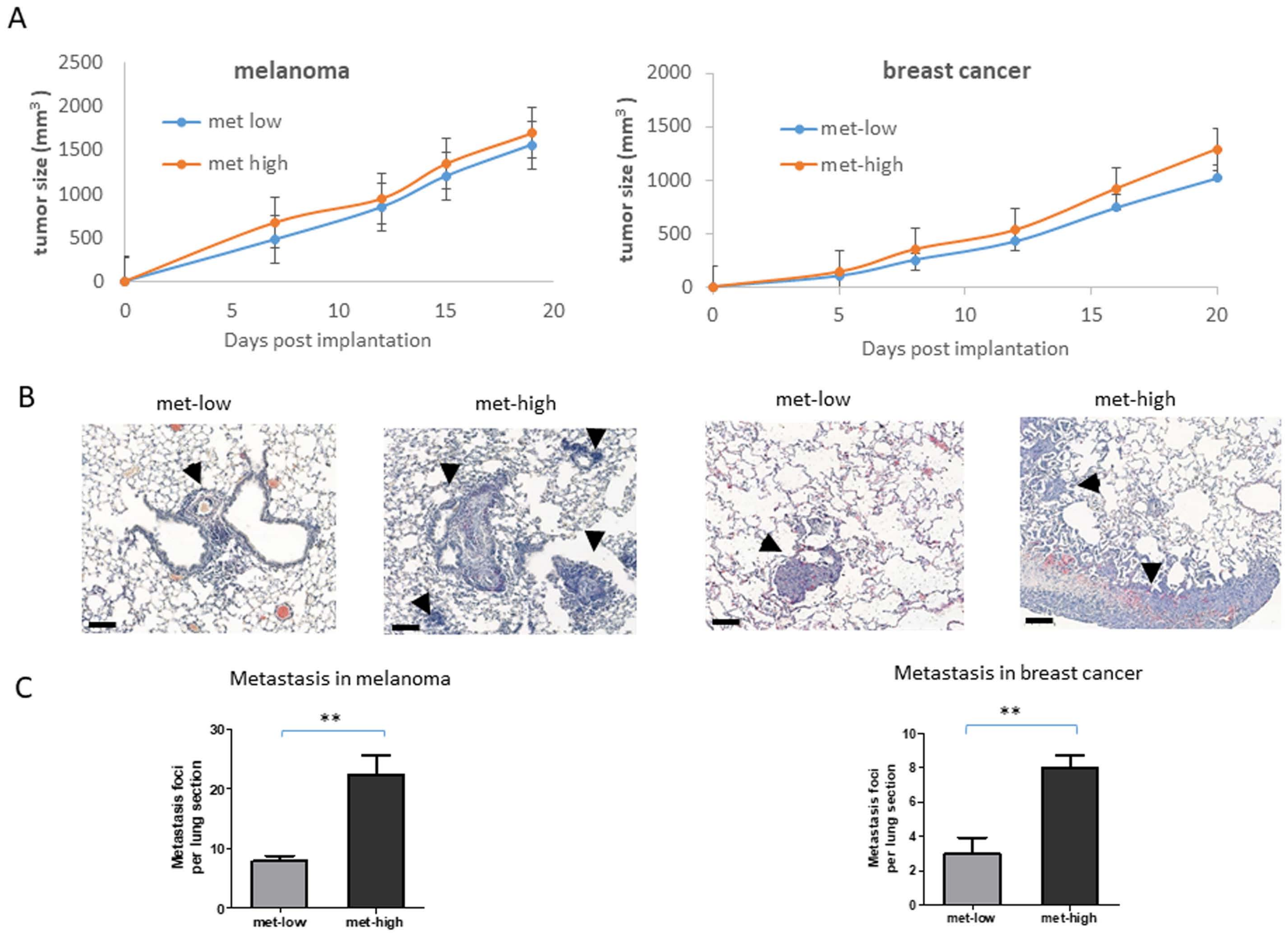
A
melanoma
tumor size (mm$^3$)
met low
met high
Days post implantation
breast cancer
met-low
met-high
B
met-low
met-high
met-low
met-high
C
Metastasis in melanoma
Metastasis foci per lung section
met-low
met-high
Metastasis in breast cancer
**

Fig. S2

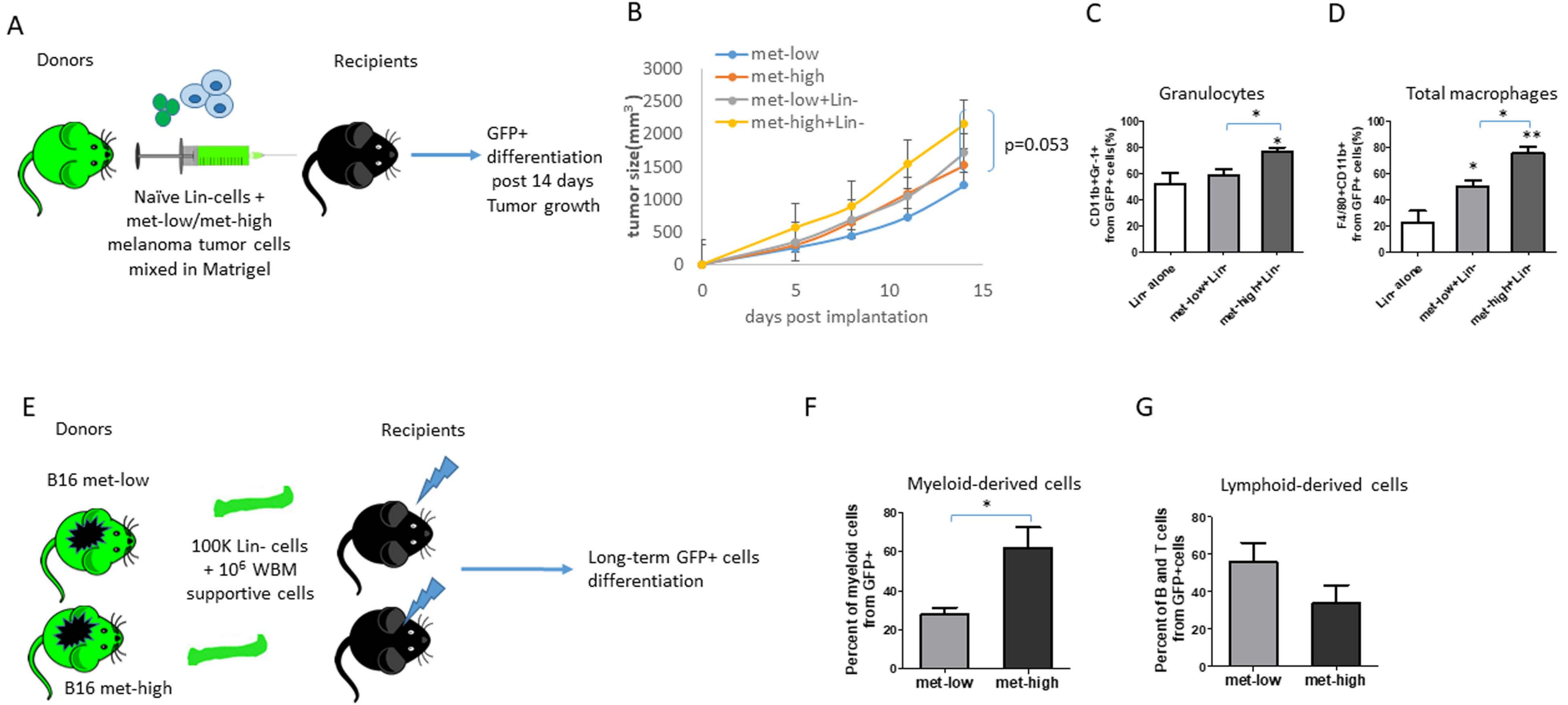
A
Donors
Recipients
Naïve Lin-cells +
met-low/met-high
melanoma tumor cells
mixed in Matrigel
GFP+
differentiation
post 14 days
Tumor growth
B
met-low
met-high
met-low+Lin-
met-high+Lin-
tumor size(mm$^3$)
3000
2500
2000
1500
1000
500
0
0
5
10
15
days post implantation
p=0.053
C
Granulocytes
CD11b+Gr-1+ from GFP+ cells(%)
100
80
60
40
20
0
Lin- alone
met-low+Lin-
met-high+Lin-
*
*
D
Total macrophages
F4/80+CD11b+ from GFP+ cells(%)
100
80
60
40
20
0
Lin- alone
met-low+Lin-
met-high+Lin-
*
*
**
E
Donors
Recipients
B16 met-low
B16 met-high
100K Lin- cells
+ 10$^6$ WBM
supportive cells
Long-term GFP+ cells
differentiation
F
Myeloid-derived cells
Percent of myeloid cells from GFP+
80
60
40
20
0
met-low
met-high
*
G
Lymphoid-derived cells
Percent of B and T cells from GFP+cells
80
60
40
20
0
met-low
met-high

Fig. S3

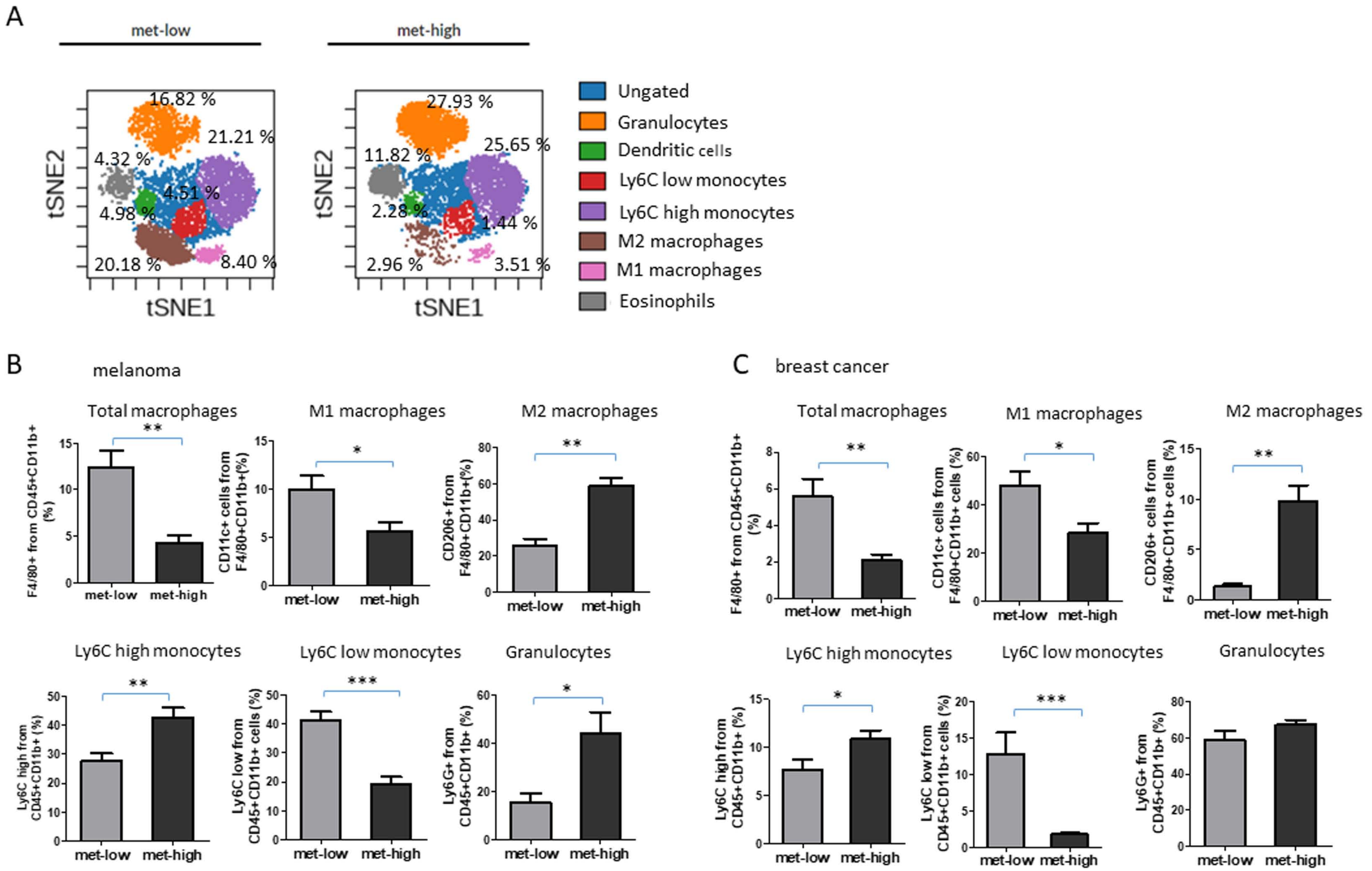

A
met-low
met-high
16.82 %
21.21 %
4.32 %
4.51 %
4.98 %
20.18 %
8.40 %
27.93 %
25.65 %
11.82 %
2.28 %
1.44 %
2.96 %
3.51 %
tSNE1
tSNE2
Ungated
Granulocytes
Dendritic cells
Ly6C low monocytes
Ly6C high monocytes
M2 macrophages
M1 macrophages
Eosinophils
B
melanoma
Total macrophages
M1 macrophages
M2 macrophages
F4/80+ from CD45+CD11b+ (%)
CD11c+ cells from F4/80+CD11b+(%)
CD206+ from F4/80+CD11b+(%)
Ly6C high monocytes
Ly6C low monocytes
Granulocytes
Ly6C high from CD45+CD11b+ (%)
Ly6C low from CD45+CD11b+ cells (%)
Ly6G+ from CD45+CD11b+ (%)
met-low
met-high
C
breast cancer
Total macrophages
M1 macrophages
M2 macrophages
F4/80+ from CD45+CD11b+ (%)
CD11c+ cells from F4/80+CD11b+ cells (%)
CD206+ cells from F4/80+CD11b+ cells (%)
Ly6C high monocytes
Ly6C low monocytes
Granulocytes
Ly6C high from CD45+CD11b+ (%)
Ly6C low from CD45+CD11b+ cells (%)
Ly6G+ from CD45+CD11b+ (%)

Fig. S4

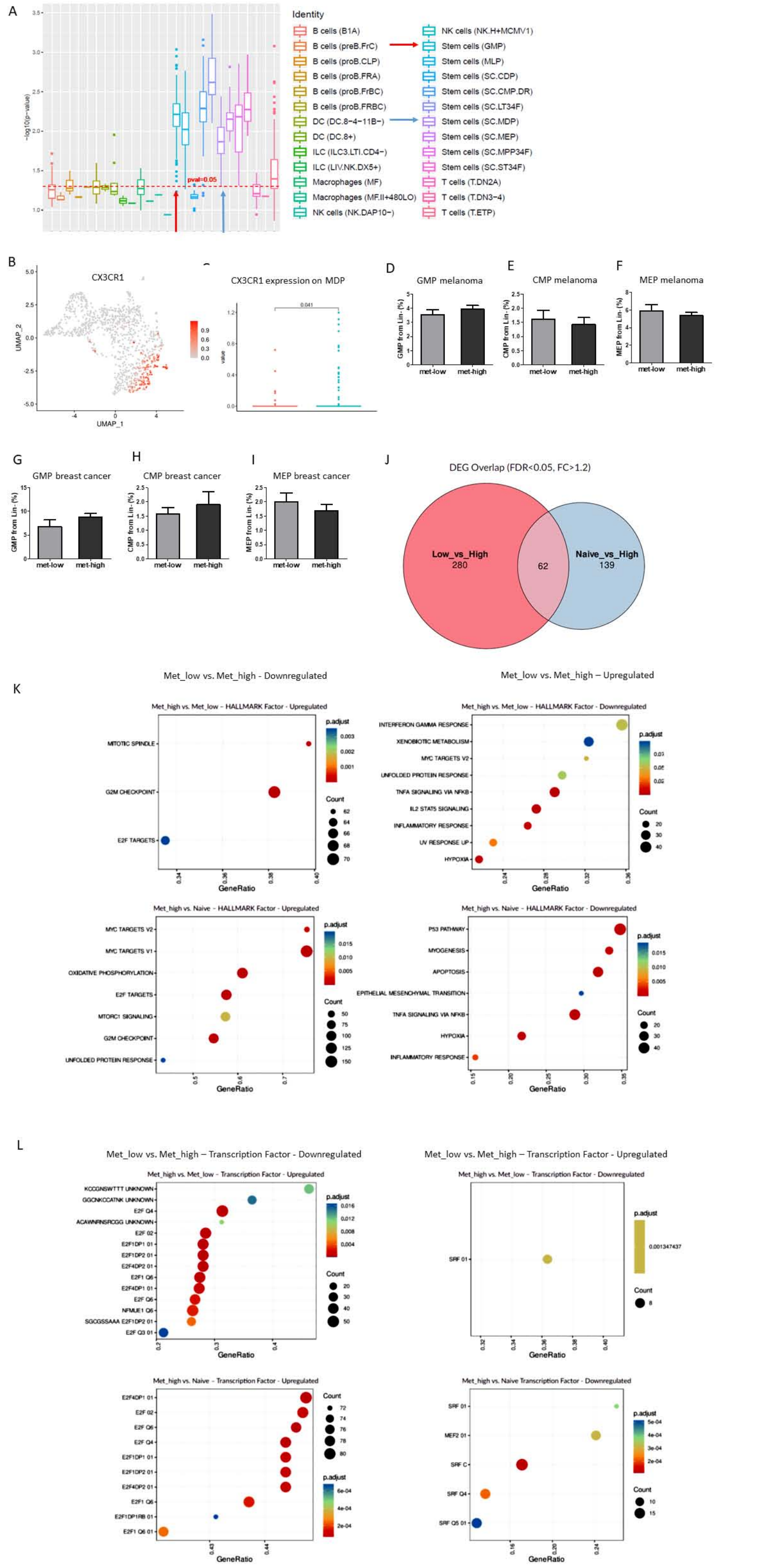
A
Identity
B cells (B1A)
B cells (preB.FrC)
B cells (proB.CLP)
B cells (proB.FRA)
B cells (proB.FrBC)
B cells (proB.FRBC)
DC (DC.8−4−11B−)
DC (DC.8+)
ILC (ILC3.LTI.CD4−)
ILC (LIV.NK.DX5+)
Macrophages (MF)
Macrophages (MF.II+480LO)
NK cells (NK.DAP10−)
NK cells (NK.H+MCMV1)
Stem cells (GMP)
Stem cells (MLP)
Stem cells (SC.CDP)
Stem cells (SC.CMP.DR)
Stem cells (SC.LT34F)
Stem cells (SC.MDP)
Stem cells (SC.MEP)
Stem cells (SC.MPP34F)
Stem cells (SC.ST34F)
T cells (T.DN2A)
T cells (T.DN3−4)
T cells (T.ETP)
pval=0.05
−log10(p−value)
B
CX3CR1
UMAP_1
UMAP_2
0.041
value
D
GMP melanoma
GMP from Lin- (%)
met-low
met-high
E
CMP melanoma
CMP from Lin- (%)
F
MEP melanoma
MEP from Lin- (%)
G
GMP breast cancer
H
CMP breast cancer
I
MEP breast cancer
J
DEG Overlap (FDR<0.05, FC>1.2)
Low_vs_High
280
62
Naive_vs_High
139
K
Met_low vs. Met_high - Downregulated
Met_low vs. Met_high – Upregulated
Met_high vs. Met_low – HALLMARK Factor - Upregulated
MITOTIC SPINDLE
G2M CHECKPOINT
E2F TARGETS
Met_high vs. Met_low – HALLMARK Factor - Downregulated
INTERFERON GAMMA RESPONSE
XENOBIOTIC METABOLISM
MYC TARGETS V2
UNFOLDED PROTEIN RESPONSE
TNFA SIGNALING VIA NFKB
IL2 STAT5 SIGNALING
INFLAMMATORY RESPONSE
UV RESPONSE UP
HYPOXIA
Met_high vs. Naive – HALLMARK Factor - Upregulated
MYC TARGETS V2
MYC TARGETS V1
OXIDATIVE PHOSPHORYLATION
E2F TARGETS
MTORC1 SIGNALING
G2M CHECKPOINT
UNFOLDED PROTEIN RESPONSE
Met_high vs. Naive – HALLMARK Factor - Downregulated
P53 PATHWAY
MYOGENESIS
APOPTOSIS
EPITHELIAL MESENCHYMAL TRANSITION
TNFA SIGNALING VIA NFKB
HYPOXIA
INFLAMMATORY RESPONSE
GeneRatio
p.adjust
Count
L
Met_low vs. Met_high – Transcription Factor - Downregulated
Met_low vs. Met_high – Transcription Factor - Upregulated
Met_high vs. Met_low - Transcription Factor - Upregulated
KCCGNSWTTT UNKNOWN
GGCNKCCATNK UNKNOWN
E2F Q4
ACAWNRNSRCGG UNKNOWN
E2F 02
E2F1DP1 01
E2F1DP2 01
E2F4DP2 01
E2F1 Q6
E2F4DP1 01
E2F Q6
NFMUE1 Q6
SGCGSSAAA E2F1DP2 01
E2F Q3 01
Met_high vs. Met_low - Transcription Factor - Downregulated
SRF 01
0.001347437
Met_high vs. Naive - Transcription Factor - Upregulated
E2F4DP1 01
E2F 02
E2F Q6
E2F Q4
E2F1DP1 01
E2F1DP2 01
E2F4DP2 01
E2F1 Q6
E2F1DP1RB 01
E2F1 Q6 01
Met_high vs. Naive Transcription Factor - Downregulated
SRF 01
MEF2 01
SRF C
SRF Q4
SRF Q5 01

Fig. S5

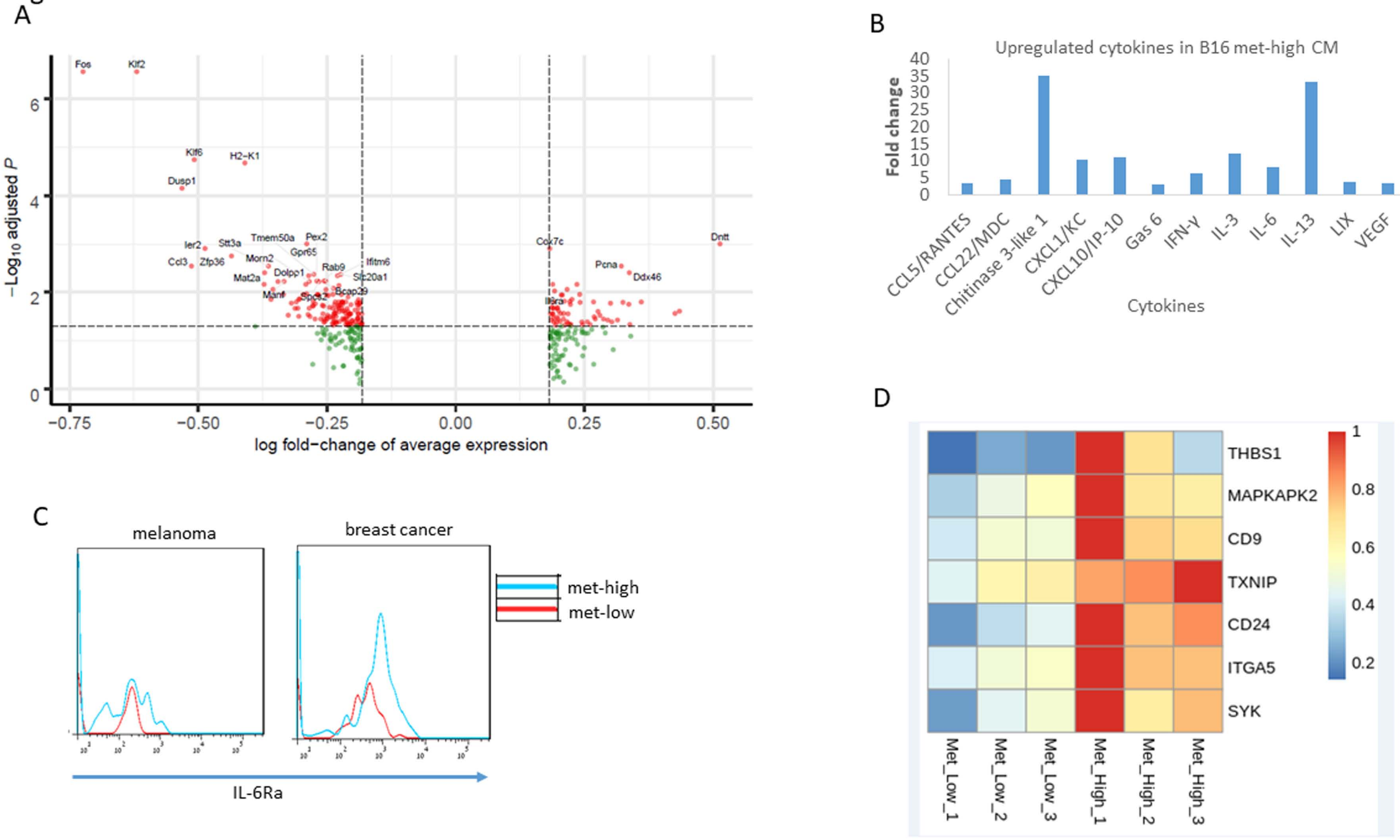

A
−Log10 adjusted P
log fold−change of average expression
Fos
Klf2
Klf6
H2−K1
Dusp1
Ier2
Stt3a
Tmem50a
Pex2
Gpr65
Morn2
Ccl3
Zfp36
Mat2a
Dolpp1
Rab9
Ifitm6
Slc20a1
Bcap29
Manf
Cox7c
Dntt
Pcna
Ddx46
B
Upregulated cytokines in B16 met-high CM
Fold change
Cytokines
CCL5/RANTES
CCL22/MDC
Chitinase 3-like 1
CXCL1/KC
CXCL10/IP-10
Gas 6
IFN-γ
IL-3
IL-6
IL-13
LIX
VEGF
C
melanoma
breast cancer
met-high
met-low
IL-6Ra
D
THBS1
MAPKAPK2
CD9
TXNIP
CD24
ITGA5
SYK
Met_Low_1
Met_Low_2
Met_Low_3
Met_High_1
Met_High_2
Met_High_3

Fig. S6

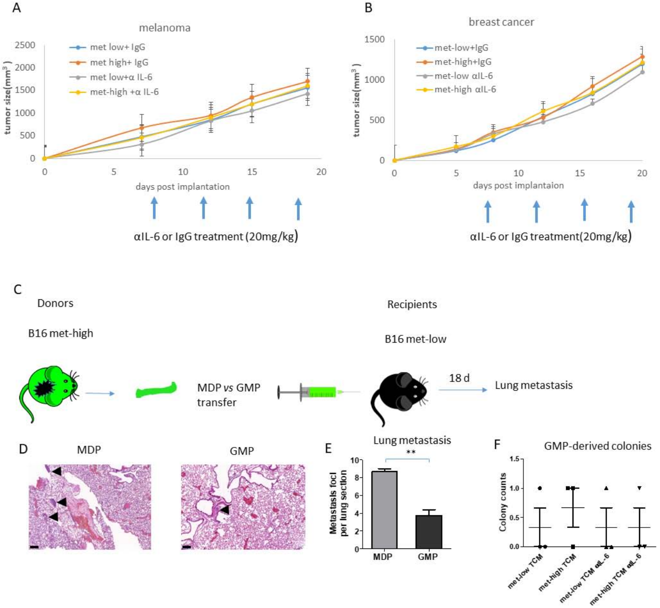
A
melanoma
met low+ IgG
met high+ IgG
met low+α IL-6
met-high +α IL-6
tumor size(mm3)
days post implantation
αIL-6 or IgG treatment (20mg/kg)
B
breast cancer
met-low+IgG
met-high+IgG
met-low αIL-6
met-high αIL-6
tumor size(mm3)
days post implantaion
αIL-6 or IgG treatment(20mg/kg)
C
Donors
B16 met-high
MDP vs GMP transfer
Recipients
B16 met-low
18 d
Lung metastasis
D
MDP
GMP
E
Lung metastasis
Metastasis foci per lung section
MDP
GMP
F
GMP-derived colonies
Colony counts
met-low TCM
met-high TCM
met-low TCM αIL-6
met-high TCM αIL-6

Fig. S7

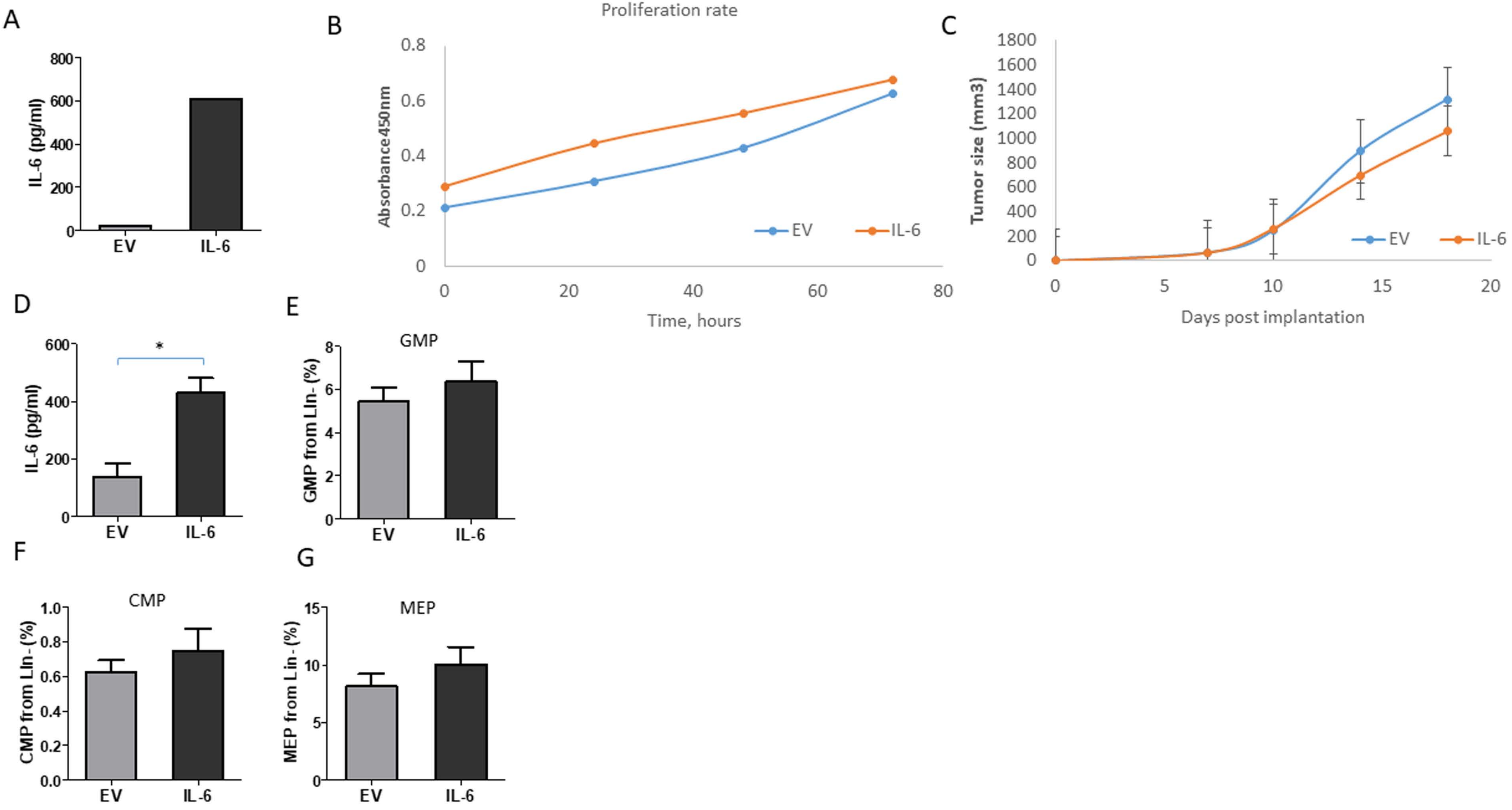

A
IL-6 (pg/ml)
800
600
400
200
0
EV
IL-6
B
Proliferation rate
Absorbance450nm
0.8
0.6
0.4
0.2
0
0
20
40
60
80
Time, hours
EV
IL-6
C
Tumor size (mm3)
1800
1600
1400
1200
1000
800
600
400
200
0
0
5
10
15
20
Days post implantation
EV
IL-6
D
IL-6 (pg/ml)
600
400
200
0
*
EV
IL-6
E
GMP
GMP from Lin- (%)
8
6
4
2
0
EV
IL-6
F
CMP
CMP from Lin- (%)
1.0
0.8
0.6
0.4
0.2
0.0
EV
IL-6
G
MEP
MEP from Lin- (%)
15
10
5
0
EV
IL-6
